# More than a prickly morphology: plastome variation in the prickly pear cacti (Opuntieae)

**DOI:** 10.1101/2023.03.13.532486

**Authors:** Matias Köhler, Marcelo Reginato, Jian-Jun Jin, Lucas C. Majure

## Abstract

Plastid genomes (plastomes) have long been recognized as highly conserved in their overall structure, size, gene arrangement and content among land plants. However, recent studies have shown that some lineages present unusual variations in some of these features. Members of the cactus family are one of these lineages, with distinct plastome structures reported across disparate lineages including gene losses, inversions, boundary movements, or loss of the canonical inverted repeat (IR) region. Here, we further investigated plastome features of the tribe Opuntieae, the remarkable prickly pear cacti, which represent a diverse and important lineage of Cactaceae. We assembled the plastome of 43 species, representing a comprehensive sampling of the tribe including all seven genera. Plastomes varied considerably in length from 121 kbp to 162 kbp, with striking differences in the content and size of the IR region (contraction and expansion events), including the lack of the canonical IR in some lineages, and the pseudogenization or loss of some genes. Overall, nine different types of plastomes were reported deviating in the presence of the IR region or the genes contained in the IR. Plastomes sequences resolved phylogenetic relationships within major clades of Opuntieae but presented some contentious nodes depending on the data set analyzed (e.g., whole plastome vs. genes only). Incongruence analyses revealed that few plastome regions are supporting the most likely topology, while disputing topologies are driven by a handful of plastome markers, which may be the result of hard recalcitrant nodes in the phylogeny or by the lack of phylogenetic signal in certain markers. Our study reveals a dynamic nature of plastome evolution across closely related lineages, shedding light on peculiar features of cactus plastomes. Variation of plastome types across Opuntieae is remarkable in size, structure, and content, and can be important for the recognition of species in some major clades. Unraveling connections between the causes of plastome variation and the consequences on species biology, ecology, diversification, and adaptation, is a promising endeavor.

## 1. INTRODUCTION

Plastids are fundamental components of plants, acting as pluripotent organelles capable of interconversion between different types - such as chromoplasts, amyloplasts, chloroplasts, etc. - for distinct functions (e.g. storage, growth, photosynthesis) (Sadali *et al*., 2019). They represent an ongoing evolutionary trajectory of endosymbiosis from a free-living prokaryote to an organelle of a eukaryotic cell, retaining the bulk of their prokaryotic biochemistry but shrunken by orders of magnitude from the genome size that their ancestors possessed (Timmis *et al*., 2004). Harboring one of the three genomes in plants, plastids are by far the most utilized for the investigation of the evolutionary history of plants, as well as physiological and adaptative features (Daniell *et al* 2016; Gitzendanner *et al*. 2018; Ruhlman & Jansen, 2021).

Within land plants, plastid genomes (plastomes) are traditionally reported as highly conserved in structure, content, arrangement, and size (Raubeson & Jansen, 2005; Wicke *et al*., 2011). Taking angiosperms as a reference, plastomes are represented as circular quadripartite monomers, highly gene dense (ca. ∼80 protein-coding genes), containing two regions of single copy (SC) distinguished by their length – a large SC (LSC, ∼80 kb) and a small SC (SSC, ∼20 kb) – separated by a large inverted repeat region (IR, approximately 25 kb) (Ruhlman & Jansen, 2021). However, advances facilitating the generation and the assembly of high-throughput molecular data of several non-model groups have increasingly reported a salient fraction of variation in plastomes (e.g. Ruhlman & Jansen, 2018; Sinn *et al*., 2018; Cauz-Santos *et al*., 2020). In this way, numerous cases of structural rearrangements including inversions and translocations (e.g., Lin *et al*. 2015; Li *et al*., 2016; Rabah *et al*., 2019; Cauz-Santos *et al*. 2020; Charboneau *et al*., 2021), pseudogenization or gene losses (e.g., Kim & Chase, 2017; Xu & Wang, 2021), as well as the lack of the IR region (e.g., Jin *et al*. 2020a; Lee *et al*., 2021) have been documented across disparate lineages, suggesting a more dynamic nature of plastome evolution, which had previously been underappreciated.

Members of the cactus family (Cactaceae, ca. 1,800 spp., Korotkova *et al*., 2021) are broadly known by their peculiar features, such as succulence, morphological diversity, spines, and exuberant flowers, making them one of the most charismatic groups of plants known worldwide (Anderson, 2001). Nonetheless, beyond their morphological, physiological, and ecological aspects, molecular components also may reveal intriguing traits of cacti. The study of entire cactus plastomes was initiated recently (Sanderson *et al*., 2015), and since then, it has been revealed that plastomes across cacti have undergone significant changes in gene content, order, and structure compared to canonical angiosperm references (Majure *et al*., 2019; Solórzano *et al*., 2019; Köhler *et al*., 2020; Oulo *et al*. 2020; Amaral *et al*., 2021; Silva *et al*., 2021; Almeida *et al*., 2021; Dalla Costa *et al*. 2022; Qin *et al*. 2022; Yu *et al*., 2023). Cactaceae seem to have the smallest plastome for an obligately photosynthetic angiosperm (∼104-113 kb, Sanderson *et al*., 2015; Solórzano *et al*., 2019; Amaral *et al*., 2021). Further, independent losses of the inverted repeat and the *ndh* gene suite have been reported in unrelated lineages (e.g., the saguaro cactus, *Carnegiea gigantea,* the cardo-ananá, *Cereus fernambucensis*, and the Chacoan-leafy-cactus, *Quiabentia verticilata,* Sanderson *et al*., 2015; Köhler *et al*., 2020; Amaral *et al*., 2021). Additionally, nearly all studied species have shown distinct features involving expansion or contractions of the IR region, rearrangements, and gene losses or pseudogenization (Solórzano *et al*., 2019; Oulo *et al*. 2020; Amaral *et al*. 2021; Silva *et al*., 2021; Almeida *et al*., 2021; Dalla Costa *et al*. 2022; Qin *et al*. 2022). Nonetheless, just a small fraction of cactus diversity (< 5%) has had their plastome analyzed in a comparative framework.

In this work, we provide a deep analysis of plastome characteristics of the tribe Opuntieae, a lineage with one of the most species-rich genera in Cactaceae (Korotkova *et al*., 2021), the emblematic prickly pear cacti (*Opuntia* spp.). The group represents a remarkable radiation of cacti broadly distributed across the major arid and semi-arid regions of the Americas (Majure *et al*., 2012; Majure & Puente, 2014), which still lack phylogenomic information to help elucidate their diversification history. We de novo assembled the plastome a comprehensive taxon sampling of major groups of the tribe, and investigate the evolution of their characteristics in a phylogenetic framework. Besides describing major features and discussing the plastome evolution in the tribe, we also investigate conflicting phylogenetic signals along different data sets (full plastome sequence vs. just plastidial genes), and performed analyses of incongruence to further assess the aspects underlying the alternative topologies.

## 2. METHODS

### 2.1. Taxon sampling, DNA extraction, and sequencing

We sampled all seven genera currently recognized and accepted within the tribe Opuntieae (Köhler *et al*., 2020; Korotkova *et al*. 2021), covering a comprehensive diversity of each genus (ranging from 15 to 100% representation), totalizing 43 accessions (Table S2). Samples are from field-collected materials or individuals grown at the Desert Botanical Garden (Phoenix, AZ, USA). DNA was extracted from epidermal tissue dried in silica gel using a modified CTAB method (Doyle & Doyle, 1987) followed by chloroform/isoamyl alcohol precipitation and silica column-based purification steps (see details in Majure *et al*. 2019). DNA quality was tested using a 1% agarose gel, and whole genomic DNAs were quantified using the Qubit dsDNA BR Assay Kit and Qubit 2.0 Fluorometer (Life Technologies, Carlsbad, CA, United States). Samples with high-molecular-weight DNA (>15 kb) showing no degradation were considered suitable and sent to Rapid Genomics LLC (Gainesville, FL, USA) for library preparation and sequencing using a genome skimming approach (Straub *et al*., 2012) on the Illumina HiSeq X platform with 150 bp paired-end reads.

### 2.2. *De novo* assemblies and annotation

Raw reads were quality controlled using BBDuk (Bushnell, 2016), removing low quality bases (Q < 20). Then, quality-controlled paired-end reads were filtered and assembled into complete plastomes using the script “get_organelle_from_reads.py” from GetOrganelle v.1.7.5 (Jin *et al*., 2020b), which uses Bowtie2 (Langmead & Salzberg, 2012), BLAST (Camacho *et al*., 2009), SPAdes 3.1.0 (Bankevich *et al*., 2012) as well as Python dependencies implemented on the HiPerGator SLURM supercomputing cluster housed at the University of Florida, Gainesville, Florida, USA. We used default settings, with kmers (-k) set as 21,45,65,85,105,115,127. When necessary, additional parameters were set (e.g., reducing word size (-w), increasing the maximum extension rounds (-R), or providing a close-relative seed database (-s)) to assemble complete graphs. Final assembly graphs were checked in Bandage (Wick *et al*., 2015) to visually evaluate their overall structures and repeated regions. Putative isomers resulting from the flip-flop recombination mediated by the IR or the sIR (short Inverted Repeat, sensu Jin *et al*. 2020c) were obtained, and one of them was arbitrarily selected for downstream analysis according to Walker *et al*. (2015) and the FAQ of GetOrganelle. Dubious boundaries junctions between IR and the SC regions, and the putative induced isomers were visually checked in Geneious using an *in-silico* approach using the library information of paired-end reads with a reference-mapping approach adapted from Jin *et al*. (2020a). Annotations were performed with GeSeq (Tillich *et al*., 2017), using default parameters to predict protein-coding genes by BLAT search, but adding as third-party references NCBI RefSeqs of *Arabidopsis thaliana* (NC_000932), *Spinacia oleracea* (NC_002202), *Solanum lycopersicum* (AC_000188) and *Glycine max* (NC_007942), and tRNAscan-SE v2.0.7 (Chan *et al*., 2021) was selected as a third party to annotate tRNA. All sequences were imported into Geneious 9.0.5 (Biomatters, Auckland, New Zealand), and their annotations were manually curated to adjust the boundaries of start and stop codons accordingly to the translated CDS of *A. thaliana* as a reference for each coding gene, or equivalent ORF of the sequence. Genes truncated with stop-codons within the frame length of the expected coding gene and exceedingly divergent protein translations (when comparing to *A. thaliana* as reference), were treated as pseudogenes, missing CDS annotation. Particularly to the *accD*, *ycf1*, and *ycf2*, we performed additional BLASTN searches (fragmenting the entire length region in 150-300 bp query) in the nt database – excluding Cactaceae from the records – to check for significant alignments with the respective CDS-feature, assessing putative loss or pseudogenization, as has been found previously in those genes (Köhler *et al*., 2020). Likewise, recognition and annotation of the LSC, SSC, IRs, and sIR were performed using Geneious based on the outputs from GeSeq and graphs analyzed in Bandage.

### 2.3. Data processing, comparative and phylogenetic analyses

We investigated the variation of plastomes of our taxon sampling in a phylogenetic framework. Considering that plastomes are represented as circular monomers, we arbitrarily established the 3’-end of the *trnH^GUG^*gene as the beginning of the monomer as a linear sequence for all plastomes in downstream analyses. Sequences had the second copy of IR or sIR removed and were visually investigated regarding the overall gene order. We also performed progressiveMauve algorithm in Mauve v2.3.1 (Darling *et al*., 2004; using Geneious plugin) with default settings to double-check our visual inspections. This analysis was performed twice, one using an outgroup as a reference to check rearrangements compared to canonical plastomes (we used *Portulaca oleracea* L., which has a canonical angiosperm plastome, and is one of the closest relatives of Cactaceae; see Walker *et al*., 2018; sequence from GenBank accession KY490694, Liu *et al*., 2018), and then one analysis only with Opuntieae samples. We categorized the Opuntieae plastomes into different types (arbitrarily named numerically following phylogenetic relationships) based on the pattern of the IR feature compared with our outgroup. We looked for Tandem Repeats in plastomes with Phobos v.3.3.12 (Mayer, 2006) using a perfect search mode, and a minimum repeat unit length of 1 bp and a maximum of 1 kb with default score constraints. We used the phylogenetic tree with more parsimony-informative sites and bootstrap means to manually map the plastome features of the lineages observed in our study (see details below).

By checking the colinear arrangement of genes within Opuntieae, we performed multiple sequence alignments using MAFFT v.7.308 (Katoh & Standley, 2013) with an automatic search for algorithm selection strategy and the default setting for score matrix and open gap penalty. We built two datasets for independent and comparative analyses, one representing the entire plastome sequences (full cpDNA) and the other including only plastidial genes (cpGenes, with coding and noncoding sequences extracted from the annotated plastome, stripping IR/sIR regions). For each of these datasets we performed four different alignment strategies for tree inference: 1) raw, as output from MAFFT; and the others trimming the alignment using GBLOCKS v.0.91.1 (Castresana, 2000; Talavera & Castresana, 2007) with default settings (minimum number of sequences for a conserved position, b1=50%+1; minimum number of sequences for a flank position, b2 = 85%; maximum number of contiguous non-conserved positions, b3=8, minimum length of a block, b4 = 10), but varying in the three options of gap presences: 2) Gblocks, no gaps allowed, 3) Gblocks, half gaps allowed, and the 4) with all gaps allowed. We then performed phylogenetic inference using Maximum Likelihood criteria implemented in RAxML 8.2.4 (Stamatakis, 2014) in the CIPRES Portal (Miller *et al*., 2010). As RAxML is mainly designed to implement generalized time-reversible molecular models (GTR), we employed the GTR + G model for the entire sequence, which has been suggested for topological reconstruction skipping model selection (Abadi *et al*., 2019), and GTR + I + G is not recommended by Stamatakis (see RAxML v8.2 manual) given the potential interaction between the I and G parameters. Support values were estimated by implementing 1,000 bootstrap pseudoreplicates, and the clade names and circumscriptions were derived from previous studies (Majure *et al*., 2012; Majure & Puente, 2014) and other ongoing projects (Majure *et al*., unpublished results).

We assessed putative phylogenetic incongruences among datasets and alignment strategies first visually. By checking incongruences between the relationship of some major clades across our datasets, we performed additional analyses, implementing the framework presented by Smith *et al*. (2015) and Shen *et al*. (2017). Briefly, we annotated intergenic spacers from the plastomes using a customized R script with functions of the genbankr package (Becker & Lawrence, 2022; all codes are available at our githubs), extracted genes and spacers, and estimated individual gene trees in RAxML as previously described. We then assessed with PhyParts (Smith *et al*., 2015) the number of markers (genes and spacers) supporting each bipartition in our primary topology (full cpDNA, raw - which yielded the highest bootstrap means and lower standard deviations values, see Results), as well as the number of markers supporting other main alternate topology, remaining alternate topologies and the number of genes not supporting any topology (neutral). PhyParts input included the primary topology (full cpDNA, raw) and gene trees for each marker, under the “fullconcon” analysis (-a 1). PhyParts output was summarized with the Python function phypartspiecharts.py (Johnson, 2017) and functions of the R package ape v.5.6.2 (Paradis & Schliep, 2019). Additionally, two major incongruent node relationships involving the positioning of BT and MSA clades (test 1), and Nopalea and Basilares clades (test 2) were investigated (see Results for details regarding topological differences). For each test, two phylogenies where only the position of these clades varied were selected from our pool of trees. Then, Shimoidara-Hasegawa tests (SH-test; Shimoidara & Hasegawa, 1999) were performed on the two alternative topologies using each individual marker alignment (genes and spacers), including raw alignments and filtered using Gblocks with no gaps allowed. The marker-wise delta log-likelihood was recorded for each comparison and significance was assessed through 10,000 bootstrap replicates. The SH-test was performed with the R package phangorn v.2.10 (Schliep 2011).

## 3. RESULTS

3.1. **Sequencing and basic assembly results –** We sequenced 43 new accessions across Opuntieae, representing all seven genera of the tribe, and a comprehensive diversity of each genus (see Methods and Tab. S1 for full details). Runs on Illumina HiSeq X resulted in 484,873,592 reads in total, between 5,279,118 (*Tacinga saxatilis*) and 19,016,846 (*Opuntia macrocentra*) per sample, for a mean read number of 11,276,130 sequences/sample. Reads per sample following quality control were between 4,881,904 and 18,531,942 with a mean post-quality control read pool number of 10,989,653 sequences/sample. The GC content following quality control were between 36.6% (*T. saxatilis*) and 40.2% (*O. austrina*). For all samples used here, we successfully assembled complete plastomes with an average base coverage varying between 80.1× (*Brasiliopuntia schulzii*) to 687.1× (*O. cuija*), and the percent of reads used for plastid assembly were between 1.08% (*B. schulzii*) and 11.57% (*O. polyacantha*).

### 3.2. Plastome features and variation within Opuntieae

Plastid genomes showed striking variation in size, varying in length from ca. 121 kb (such as in *Tacinga saxatilis* and *Opuntia polyacantha*) to ca. 160-162 kb (e.g., *Brasiliopuntia* spp. and *Consolea* spp.) (Fig. 1, Tab. 1). This size variation was remarkably associated to the dynamic movement of expansion or contraction of the gene content in the inverted repeat region across Opuntieae, including the lack of the canonical IR in *Tacinga* spp. and in members of the Basilares, Scheerianeae, Setispina, Macrocentra, and Humifusa clades – which have atypical short-IR regions (sIR, varying from 183 bp to ca. 2 kb), and an exceptional case of a short direct repeat (sDR, 793 bp) and a pair of sIR (657 and 708 bp) in *O. polyacantha* (Fig. 1 and S1; Tab. 1).

**Figure 1.**
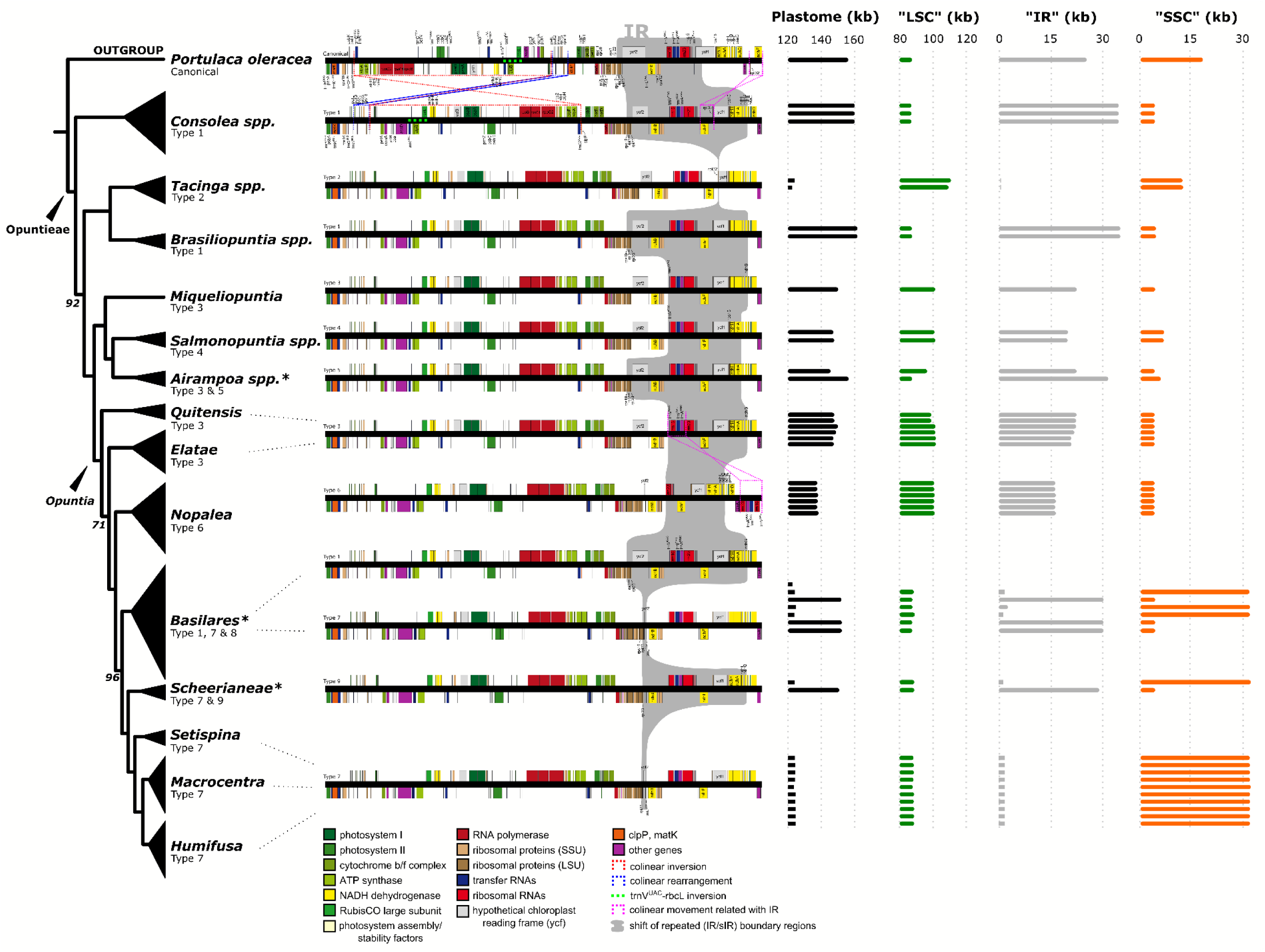
Plastome variation across Opuntieae lineages. The linear sequence of the plastomes are depicted accompanying the phylogenetic relationships inferred for the tribe (for plastome type 8, see Fig. S1). On the right, bar plots indicate variation in the size of plastome and their content. Variations in the IR-feature of plastome types are highlighted by the gray box in the linear plastome sequences. On the tree tips, asterisks (*) indicate that more than one plastome type is recovered in the clade. On the bar plot titles, quotes (“) indicate that the term is not fully coherent with some plastomes considering that they lack an IR, so the use of LSC or SSC is just for convenience across the dataset. Phylogenetic nodes have total bootstrap support values, except when noted.

**Table 1.**
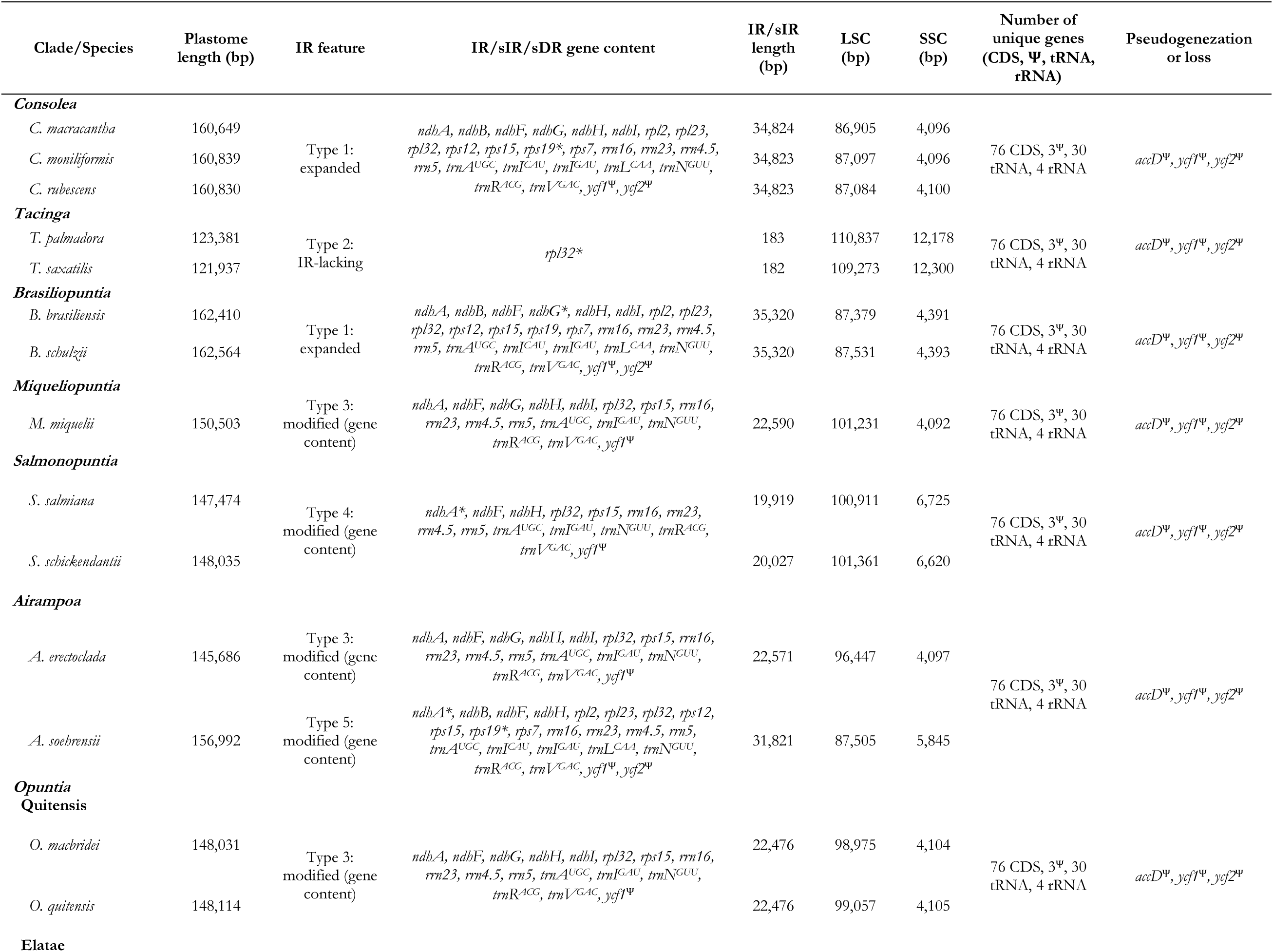
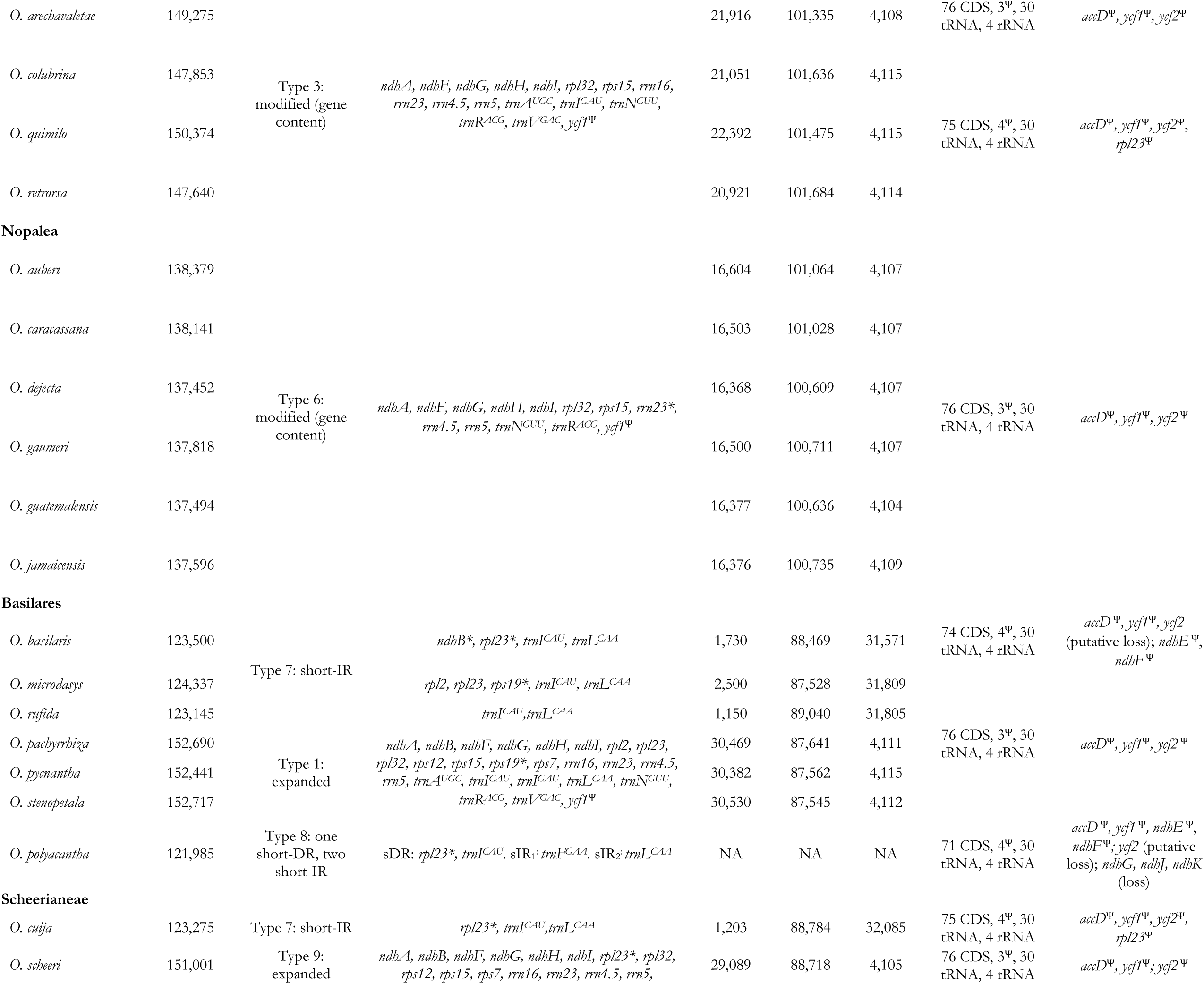
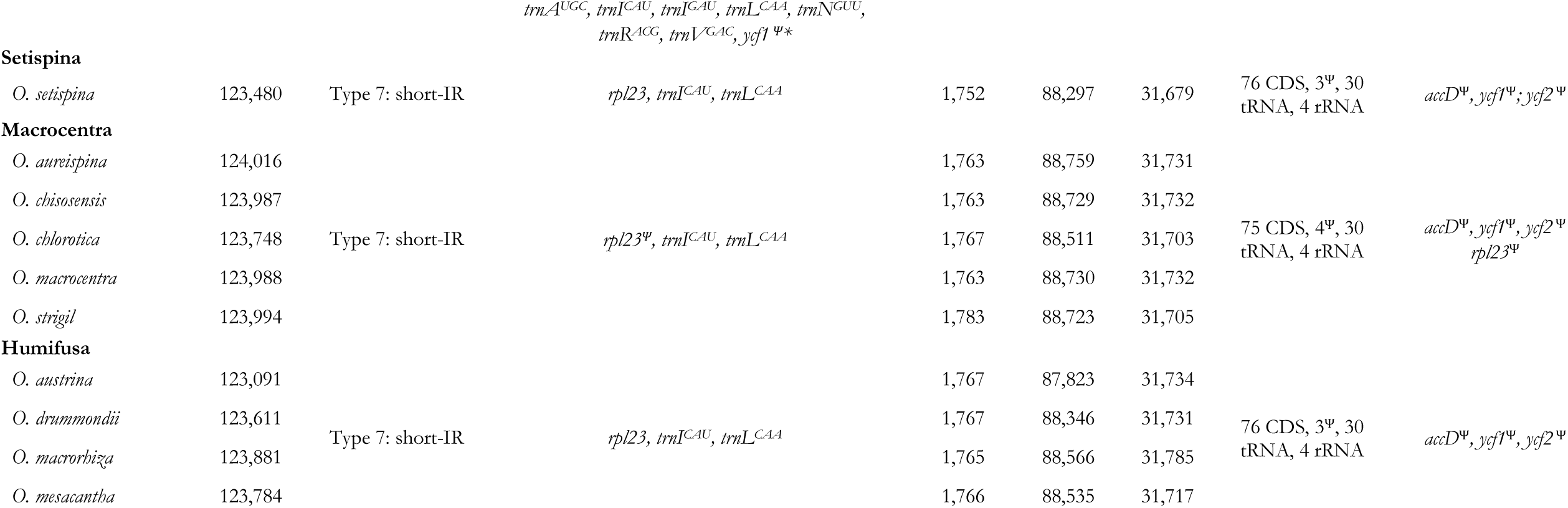
Plastome features of Opuntieae members. Asterisks (*) indicate that the gene is located in the boundary of IR and SC, being fragmented in one of the IR/SC junctions. Psi (Ѱ) indicates pseudogene.

In total, nine distinct types of plastomes were assembled and annotated based on the pattern of the IR feature considering their presence or absence, gene content, and size (Fig. 1 and S1; Tab. 1). Three type of plastomes lack the canonical IR: the type 2, in *Tacinga* spp., presenting only a pseudogenized and fragmented remnant *rpl32 ^Ѱ^* of ca. 180 bp; the type 7, in members of Basilares, Scheerianeae, Setispina, Macrocentra, and Humifusa clades, which presents short-IRs of ca. 1-2kb mostly reduced to *rpl23* (eventually fragmented or pseudogenized), *trnI^CAU^*, *trnL^CAA^*, and a pseudogenized *ycf2 ^Ѱ^*; and the type 8, so far unique in *O. polyacantha*, presenting only a sDR of *rpl23* and *trnI^CAU^*, and a pair of sIRs, one of *trnF^GAA^* and other of *trnL^CAA^* (Tab. 1). The other six plastome types (Type 1, 3, 4, 5, 6, and 9) differ in their gene content and size of the IR, varying in the IR size between ca. 16 kb (Nopalea clade) to ca. 35 kb (*Consolea* and *Brasiliopuntia* clades), containing most of the typical IR genes, but with notable expansions (encompassing genes usually present in the SSC), or the transfer of typical IR-genes to the single copy region.

In general, Opuntieae plastomes contain 110 unique genes, including 76 protein coding sequences (CDS), 30 transfer RNAs (tRNA), and four ribosomal RNAs (rRNA) (Tab. 1). Variations are observed when putative pseudogenization or gene losses are presented in some plastomes, such as the *rpl23* (which is pseudogenized in some lineages with plastomes type 3 and 7; Elatae, Scheerianeae, and Macrocentra clades), and some genes of the *ndh* suite (e.g. *ndhE* and *ndhF* which are pseudogenized in *Opuntia basilaris* and *O. polyacantha;* and *ndhG, ndhJ, and ndhK* which are lost in *O. polyacantha*). Additionally, all Opuntieae plastomes seem to have undergone pseudogenization of the *ycf1, ycf2*, and *accD*; the loss of one intron in *rpl2*, and two introns in *clpP*. The *ycf1* region varied from ca. 2 kb (*O. retrorsa* and *O. colubrina*) to ca. 4 kb (*Consolea* spp.), in all cases containing fragmented ORFs within the length-frame of the expected product of the gene, accumulating highly divergent sequences and conserving just small fragments with BLAST-identity to conserved *ycf1* of other angiosperms lineages. Similarly, the *ycf2* region varied greatly across Opuntieae, from ca. 328 bp (e.g., *O. rufida*, *O. microdasys*, *O. stenopetala*) to more than 6 kb (e.g., *O. colubrina*, *O. arechavaletae*, *Brasiliopuntia* spp.), also accumulating highly divergent sequences, preserving just small fragments with BLAST-identity to conserved *ycf2* genes of other angiosperm lineages, with putative loss and a second degradation in some lineages. The *accD* region is presented as a long ORF of ca. 3.5 kb, accumulating long and divergent fragments of sequences, and conserving just a small domain (ca. 1.4 kb) with identity to canonicals *accD* of other angiosperm lineages.

All Opuntieae plastomes assembled here shared rearrangements involving two blocks, an apparent translocation of the *petL–rps12*^exon^ ^1^ (ca. 4 kb; Fig. 1, blue dotted line), and an inversion involving the *trnG^UCC^*–*psbE* region (ca. 58 kb; Fig. 1, red dotted line), compared with canonical angiosperm plastomes, as represented by *Portulaca oleracea* (Liu *et al*., 2018). Considering the adjacency of the two rearranged blocks, the rearrangement might also be caused by two inversion events, an inversion of the *trnG^UCC^–rps12*^exon^ ^1^ region followed by an inversion of the *rps12*^exon^ ^1^*– petL* region. Within the trnG^UCC^–*psbE* block, a second inversion occurred involving the *trnV^UAC^*–*rbcL* inversion (ca. 5 kb; Fig. 1, green dotted line), which turned the region of *accD* adjacent to the *trnV^UAC^*, and not to *rbcL*, as is in canonical angiosperm plastomes.

Opuntieae plastomes harbor a significant number of tandem repeats (simple sequence repeats – cpSSRs) which are relatively conserved in number among lineages, varying from 816 (*Opuntia guatemalensis*) to 857 (*Salmonopuntia schickendantzii*), totaling ca. 7.4 % to 8.4 % of the plastome content (*O. colubrina* and *Tacinga palmadora*, respectively) (Tab. 2, and Tab. S2 for full data). Most of the SSRs are mononucleotide repeats derived from A/T varying in length from 7 to 29 bp, but a remarkable presence of more complex repeats (penta, hexa, and higher than seven nucleotides) is noteworthy in number and length.

**Table 2.** Summary information of number, type, and variation in length (bp) of tandem repeats (Simple Sequence Repeats – SSRs) within Opuntieae plastomes.

| Clade | Total number of repeats | Total length of repeats (bp) | Plastome repeat content (%) | Type of repeat |  |  |  |  |  |  |  |  |  |  |  |  |  |
| --- | --- | --- | --- | --- | --- | --- | --- | --- | --- | --- | --- | --- | --- | --- | --- | --- | --- |
|  |  |  |  | Mononucleotide |  | Dinucleotide |  | Trinucleotide |  | Tetranucleotide |  | Pentanucleotide |  | Hexanucleotide |  | >7 |  |
|  |  |  |  | Total | Length | Total | Length | Total | Length | Total | Length | Total | Length | Total | Length | Total | Length |
| Consolea | 838–846 | 10,005–10,233 | 7.9–8.1 | 336–338 | 7–17 | 41 | 8–16 | 59 | 9–13 | 86–87 | 10–14 | 99–100 | 11–19 | 131–132 | 12–19 | 85–89 | 14–348 |
| Brasiliopuntia | 843–852 | 9,826–9,982 | 7.7–7.8 | 332–334 | 7–18 | 42 | 8–26 | 57 | 9–15 | 89–92 | 10–14 | 94–96 | 11–17 | 133–134 | 12–17 | 96–97 | 14–183 |
| Tacinga | 832–835 | 10,111–10,361 | 8.3–8.4 | 332–334 | 7–19 | 41–42 | 8–20 | 53 | 9–15 | 88–93 | 10–14 | 95–97 | 11–19 | 125–126 | 12–80 | 91–97 | 14–287 |
| MST | 829–857 | 9,599–9,949 | 7.5–7.8 | 336–348 | 7–25 | 40–41 | 8–25 | 57–62 | 9–13 | 87–91 | 10–14 | 93–101 | 11–133 | 125–130 | 12–126 | 85–90 | 14–220 |
| Quitensis | 850–852 | 9,820–9,923 | 7.8 | 340 | 7–17 | 44 | 8–28 | 57–58 | 9–13 | 91–92 | 10–15 | 101 | 11–19 | 126 | 12–42 | 91 | 14–168 |
| Elatae | 838–857 | 9,434–9,711 | 7.4–7.6 | 335–343 | 7–19 | 42–44 | 8–20 | 58–62 | 9–26 | 88–95 | 10–14 | 95–102 | 11–20 | 125–127 | 12–17 | 84–88 | 14–254 |
| Nopaleae | 816–824 | 9,359–9,969 | 7.7–8.1 | 327–332 | 7–26 | 40–43 | 8–22 | 56 | 9–13 | 88–91 | 10–14 | 102–103 | 11–25 | 117–119 | 12–33 | 81–86 | 14–360 |
| Basilares | 826–845 | 99,487–10,104 | 7.7–8.3 | 328–336 | 7–24 | 39–43 | 8–24 | 57–63 | 9–20 | 87–91 | 10–16 | 92–104 | 11–70 | 120–125 | 12–63 | 87–95 | 14–372 |
| Scheerianeae | 831–837 | 9,462–9,937 | 7.7–8.1 | 339–340 | 7–17 | 43 | 8–20 | 54–55 | 9–13 | 87–89 | 10–14 | 101–102 | 11–25 | 123–125 | 12–17 | 81–86 | 14–357 |
| Setispina | 829 | 9,316 | 7.6 | 336 | 7–29 | 44 | 8–26 | 55 | 9–13 | 86 | 10–14 | 99 | 11–19 | 128 | 12–17 | 81 | 14–198 |
| Humifusa | 823–834 | 9,251–9,812 | 7.6–8 | 332–334 | 7–16 | 41–42 | 8–20 | 55–57 | 9–13 | 86 | 10–14 | 97–99 | 11–19 | 126–129 | 12–19 | 83–89 | 14–191 |
| Macrocentra | 823–828 | 9,468–9,796 | 7.7–8 | 331–333 | 7–21 | 42 | 8–24 | 55–56 | 9–15 | 85–87 | 10–16 | 98–99 | 11–20 | 126–128 | 12–17 | 83–86 | 14–215 |

### 3.3. Phylogenetic relationships, and plastome evolution

Plastid genome sequences were shown to be informative to infer phylogenetic relationships within Opuntieae lineages, supporting eleven major clades based on our sampling with high support (Fig. 2, highlighted and annotated clades). The full cpDNA dataset resulted in a raw alignment of 137,064 bp, with 5,135 distinct patterns, 3,769 parsimony-informative sites (PIS), and 129,283 constant sites; whereas the cpGenes resulted in a raw alignment of 85,574 bp, 2,677 distinct patterns, 2,155 PIS, and 82,324 constant sites (results for all alignment scenarios are in Tab. S3). All full plastome alignment scenarios (raw and three Gblocks settings) recovered the same backbone topology (Fig S2), with the raw alignment yielding higher bootstrap means and lower standard variations (Tab. S3). However, contrasting topologies were recovered by comparing the full plastome dataset (full cpDNA) to the just genes (cpGenes) datasets involving two nodes (Fig. 2A-C; Fig. S2). *Consolea* was sister to the rest of Opuntieae, while the relationship of the BT (*Brasiliopuntia* + *Tacinga*) or MSA (*Miqueliopuntia* + *Salmonopuntia* + *Airampoa*) clades as sister to the *Opuntia* clade was conflicting when comparing the full cpDNA to the cpGenes datasets – despite both scenarios recovering high bootstrap support values. A similar conflicting topology was recovered within *Opuntia* lineages, with the full cpDNA and the cpGenes (raw alignment) supporting the Nopalea clade as sister to the rest of the North American (NA) *Opuntia* clades (Basilares + Scheerianeae + Setispina + Humifusa + Macrocentra), while the cpGenes (Gblocks no gaps, half gaps, and all gaps allowed) recovered a third topology, suggesting a sister-relationship between the Nopalea and Basilares clades, but with low bootstrap support (Fig. 2C).

**Figure 2.**
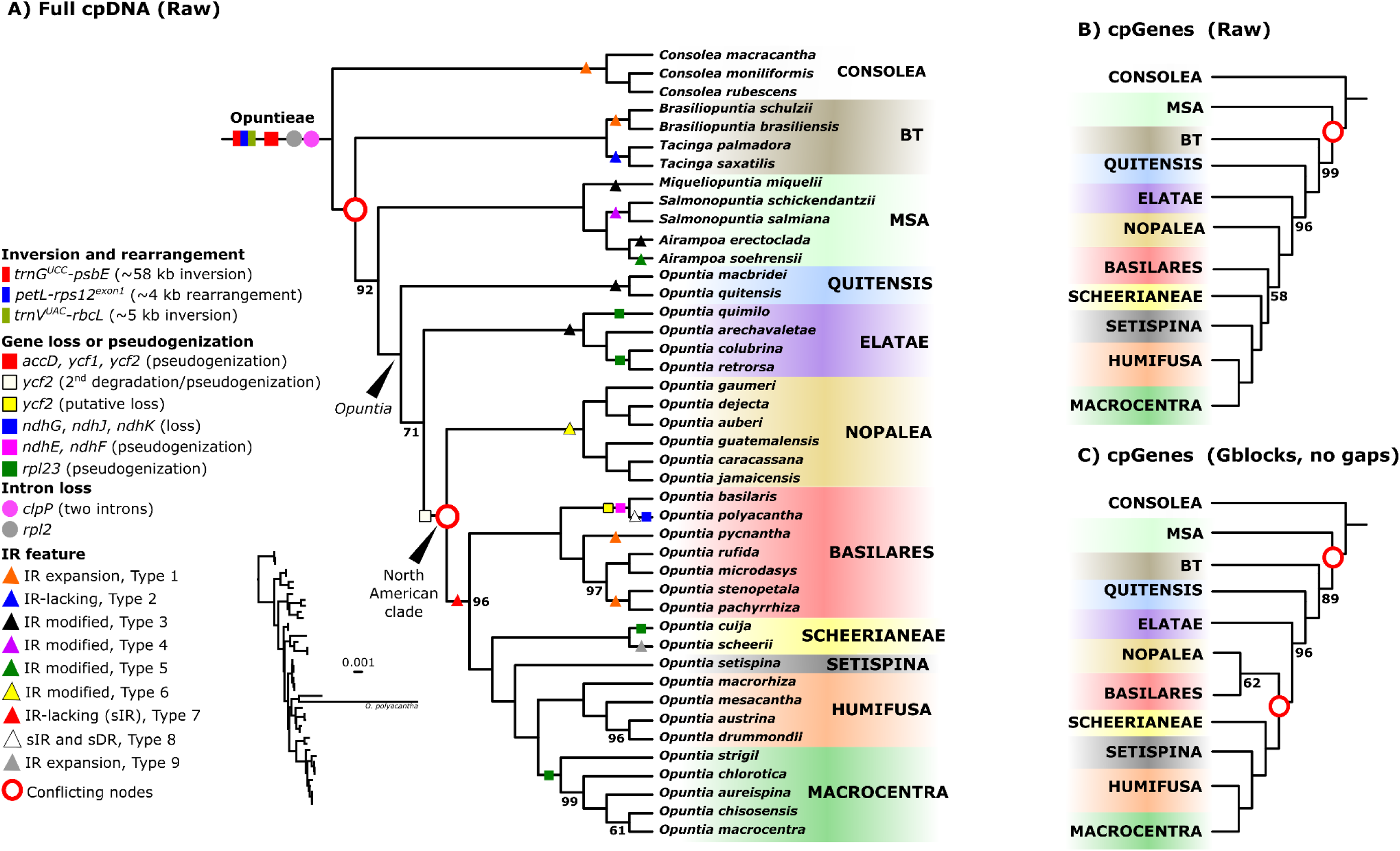
Phylogenetic inference of tribe Opuntieae based on plastome sequences with the maximum likelihood (ML) criterion, with the major plastome evolution features mapped on the tree. **A.** Inference based on the full plastome sequences (genes and intergenic regions, raw alignment). **B.** Inference based on chloroplast genes (raw alignment). **C.** Inference based on plastome genes trimmed with Gblocks (no gaps allowed). Phylogenetic nodes have full bootstrap support (100), except when depicted; incongruent nodes across different datasets are highlighted in red.

Incongruence analyses revealed that most nodes of the full cpDNA (raw)-based phylogeny are supported by a few regions of the plastome sequence (Fig. 3A), even when they have maximum bootstrap support value (bs=100; e.g., North American clade of *Opuntia* has total bootstrap value, and is supported by 15 plastome regions). Similarly, the alternative topologies regarding the two contentious relationships involving 1) the MSA or BT clade as sister to *Opuntia* (Fig. 3B), or 2) the Nopalea clade as sister to Basilares or sister to the rest of the NA *Opuntia* clades (Fig. 3C) are mainly driven by a handful of markers. However, while the MSA/BT recalcitrant relationship seems to be a hard incongruence with around a dozen markers disputing the alternative topologies, the Nopalea/Basilares contentious relationship seems to be more by a lack of resolution across the analyzed regions than a dispute for alternative topologies.

**Figure 3.**
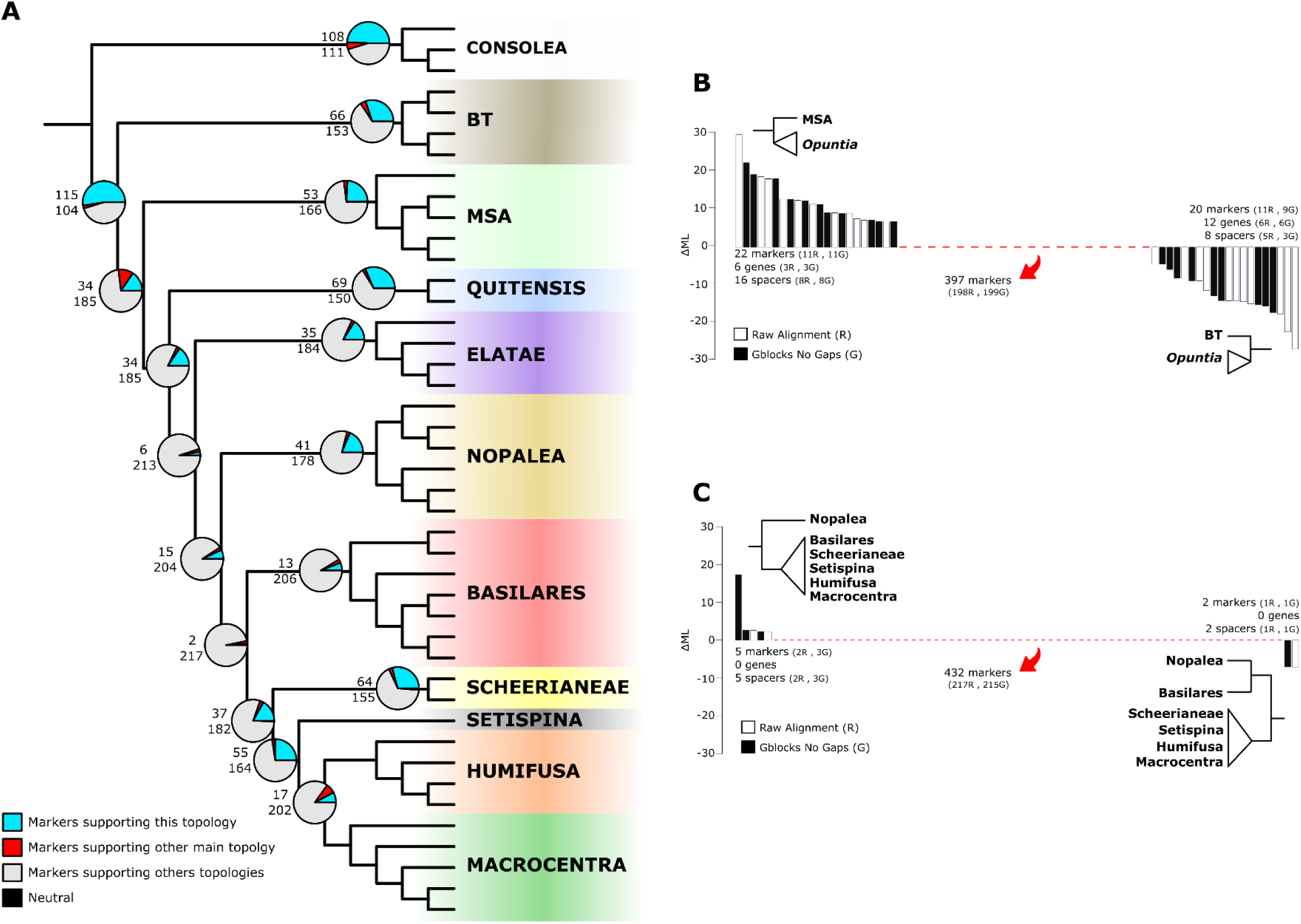
Incongruence analyses. **A.** Pie charts depicting the number of plastome markers (gene or spacers regions) supporting the shown topology (light-blue, numbers above), or supporting others topologies (number below). **B. and C.** Marker-wise delta log-likelihood (ΔML) for the alternative topologies; bar plots shown with positive values are markers significantly supporting the schematic topology on the left, while negative values are markers supporting the right one. Red bar plots are neutral markers, with no significative support for any of the compared topologies.

Regardless of incongruent topologies, *Opuntia* is well resolved as monophyletic including eight major clades (Fig. 2). The South American (SA) Quitensis clade (including *O. macbridei* and *O. quitensis*) is sister to the rest of the genus, while the other SA clade, Elatae (inc. *O. retrorsa*, *O. colubrina*, *O. arechavaletae*, and *O. quimilo*), is resolved as sister to the North American (NA) clade. The NA clade encompasses the Nopalea clade (inc. *O. dejecta*, *O. auberi*, *O. gaumeri*, *O. guatemalensis*, *O. caracassana*, and *O. jamaicensis*) as sister to the rest of the NA clades (in the full cpDNA and cpGenes raw), or as sister to the Basilares clade (inc. *O. pachyrrhiza*, *O. stenopetala*, *O. microdasys*, *O. rufida*, *O. pycnantha*, *O. polyacantha*, and *O. basilaris*) in the cpGenes (Gblocks scenarios). The Scheerianeae clade (*O. scheerii*, and *O. cuija*) and the Setispina (*O. setispina*) clades are subsequent sisters to a clade with the Humifusa (*O. drummondii*, *O. austrina*, *O. mesacantha*, *O. macrorhiza*) + Macrocentra clades (*O. macrocentra*, *O. chisosensis*, *O. aureispina*, *O. chlorotica*, *O. strigil*).

The evolution of plastome types across Opuntieae appears to be both homoplasious (presenting recurrent features in several different clades), as well as clade- or species-specific. The largest plastomes of Opuntieae (ca. 160-162 kb, plastome Type 1), which have incorporated some genes that are typically in the SSC into the IR region, are present in the *Consolea* and *Brasiliopuntia* clades but are also found in distantly related species of the Basilares clade (*Opuntia pycnantha*, *O. stenopetala*, and *O. pachyrrhiza*). Similarly, the plastome Type 3, which is ubiquitous in the Elatae clade, is also observed in the Quitensis clade and members of the MST clade. On the other hand, the plastome Type 6, which has transferred some rRNA genes that are typically in the IR region to the SC region, is exclusive of the Nopalea clade, while the IR-lacking plastome Type 2 is exclusive of the *Tacinga* clade. Likewise, the IR-lacking plastome Type 7 is the most common in the North American *Opuntia* clade (pervasive in the Macrocentra, Humifusa, and Setispina clades), although not exclusive to all lineages.

A similar pattern is observed with gene loss and some pseudogenizations. Despite the major inversions (*trnG^UCC^-psbE*, *trnV^UAC^-rbcL*), rearrangement (*petL-rps12^exon1^*), pseudogenization (*accD*, *ycf1,* and *ycf2*), and intron loss (*clpP* and *rpl2*), which were observed in all Opuntieae plastomes, other events seem to have occurred independently. The *rpl23* pseudogenization is characteristic of the Macrocentra clade but is also present in independent lineages of the Scheerianeae and Elatae clades. On the other hand, the second degradation/pseudogenization of *ycf2*, reducing the large region of ca. 3-6 kb to only ca. 400 bp, has occurred once for the NA clade. Similarly, the putative loss of *ycf2* and the *ndhE*-*ndhF* pseudogenization is only observed in the sister taxa, *Opuntia basilaris* + *O. polyacantha*, while the *ndhG*, *ndhJ,* and *ndhK* loss is unique in *O. polyacantha*.

## 4. DISCUSSION

The structure, content, and arrangement of plastomes have been largely studied in the last decades, especially increasing with the advent of the high-throughput sequencing era, which has revolutionized innumerable aspects of evolutionary biology. Despite numerous advances supporting the overall conserved structure of plastomes in land plants, there are also accumulating reports of variation in the theme across disparate lineages, which have challenged the idea of an overall conserved structure among plant lineages (Mower & Vickrey, 2018; Ruhlman & Jansen, 2018; Ruhlman & Jansen, 2021). Furthermore, other misconceptions may obfuscate our understanding of this organelle evolution (Gonçalves *et al*., 2020). For example, the very idea of the plastome as a circular molecule – which seemed intuitive as bacterial descendants through the endosymbiotic origin of the organelle – has been rethought with the understanding of predominantly linear and branched molecules covalently linked through repeating units that undergo recombination both within and between units and molecules (Bendich & Smith, 1990; Lilly *et al*., 2001; Bendich, 2004; Oldenburg and Bendich, 2004, 2015, 2016; Lee *et al*., 2021). In this scenario, the mechanisms underlying such diversity of variation in content and structure of plastomes, as reported here, can be more complex to discern, but at the same time are more likely to occur. However, the incorporation of these emerging ideas in the ongoing flux of data with robust bioinformatics tools is still a bottleneck to providing a deeper understanding of plastome evolution.

The transfer of genes between SC regions to the IR (shifts in the IR boundaries), and vice versa, is a frequent mechanism involved in plastome evolution, and it seems to have a major impact on plastome variation across disparate and closely related species (Ruhlman & Jansen, 2018; Zhu *et al*., 2016; Choi *et al*., 2020). In Opuntieae, this seems to be the main mechanism associated with the variation of plastome types observed here, accompanied by the events of loss of the IR region. Interestingly, plastomes within Opuntieae are well conserved regarding gene arrangement, conserving a colinear syntenic block of genes, which stresses a standing idea that rearrangements are more frequent when the large inverted repeat is lost (Palmer & Thompson, 1982). Although the presence of repetitive sequences in plastomes has been reported as an important mechanism in plastome size evolution and genomic rearrangements (Jo *et al*., 2011; Dugas *et al*., 2015; Wu *et al*., 2021), our data do not fully support these hypotheses, considering that Opuntieae plastomes do not present significant rearrangements. Furthermore, despite Opuntieae presenting a striking variation in plastome size across major clades, the number of tandem repeats is quite similar among the clades. However, our data may help to further assess the impact of tandem repeats in the pseudogenization and shifts in the inverted repeat region (Sinn *et al*., 2018).

On the other hand, compared to canonical angiosperms as represented by the outgroup *Portulaca oleracea*, Opuntieae plastomes have notable rearrangements involving two blocks of sequences with putative inversions and translocations. All Opuntieae samples presented the *trnV^UAC^*–*rbcL* inversion (ca. 5 kb), which has long been proposed as a synapomorphy of Cactaceae (Downie & Palmer, 1994; Wallace, 1995). Some of these inversions may likely have influenced a few features observed in some of the genes adjacent to the regions involved in the inversions, such as the *clpP* (which has lost two introns and is reduced to a small, conserved domain, ca. 150 bp), and the *accD* (which is presented as a long ORF of ca. 3.5 kb, accumulating a long and divergent fragment of sequence, and just a small conserved domain – putatively pseudogene). However, these genes are also widely reported as pseudogenes, lost, or under positive selection in several disparate lineages not necessarily related to inversions (e.g.: Jansen *et al*., 2008; Wicke *et al*., 2011; Harris *et al*., 2013; Dugas *et al*., 2015; Wang *et al*., 2016; Ruhlman & Jansen, 2018), suggesting that other mechanisms may be involved in their modifications. Also, we have reported *ycf1* and *ycf2* pseudogenization, which are not directly involved in rearrangement breakpoints but are in shift movements of SC/IR regions, especially *ycf2*. Although both genes appear to be essential for plastid function in most plants (Drescher *et al*., 2000; De Vries *et al*., 2015; Yang *et al*., 2016; Kikuchi *et al*., 2018), their loss or pseudogenization is relatively frequent (see Graham *et al*., 2017; Ruhlman & Jansen, 2021). For example, all Poales have lost the *ycf1* and *ycf2* in a progressive degradation of the gene sequence (Guisinger *et al*., 2010; de Vries *et al*., 2015), which could also have occurred in Opuntieae since the loss of *ycf2* and small degraded fragments (ca. 300 bp) are shared feature of the most derived clades of *Opuntia* (the North American *Opuntia* clade: Nopalea + Basilares + Scheerianeae + Setispina + Humifusa + Macrocentra). However, the mechanisms responsible for such aspects remain elusive.

Other events of pseudogenization, gene, and intron loss are remarkable within Opuntieae, representing unique or shared occurrences. The *rpl2* intron loss is ubiquitous in Opuntieae, and has been reported as synapomorphic within Centrospermae lineages (Caryophyllales, Palmer *et al*., 1988; Yao *et al*., 2019). Similarly, the events of loss or pseudogenization of *ndh* suite genes have been reported in some cactus lineages (Sanderson *et al*., 2015; Solórzano *et al*., 2019; Köhler *et al*., 2020; Amaral *et al*. 2021; Dalla Costa *et al*., 2022; Qin *et al*., 2022; Silva *et al*., 2021), as it is especially associated with hemi- or holopasrasitism, carnivory, xerophytes and submersed plants (Braukman *et al*., 2009; Wicke *et al*., 2011; Peredo *et al*., 2013; Silva *et al*., 2016). Previous authors have suggested that the retention of the *ndh* complex is associated with the transition of plants to stressful environments and that *ndh* loss would be associated with decreased environmental stressors but of limited biological significance in contemporary plants (Ruhlman *et al*., 2015; Lin *et al*., 2017). The association between *ndh* loss and the presence of Crassulacean Acid Metabolism (CAM) photosynthesis has also been speculated (Strand *et al*. 2019; Köhler *et al*., 2020), but is still elusive. On the other hand, a dynamic transfer of segments of the plastid genome to the nuclear or mitochondrial genome - and vice versa - has been increasingly reported (Stegemann *et al*. 2003; Cui *et al*., 2021; Hertle *et al*., 2021), which could suggest the transition of these genes to other genomes. Sanderson *et al*. (2015) found many nonplastid copies of plastid *ndh* genes in the nuclear genome of the saguaro cactus, but none had intact reading frames. Based on the presence of other nuclear genes, they conclude the existence of an alternative pathway redundant with the function of the plastid NADH dehydrogenase-like complex (*ndh*) may have facilitated the loss of the plastid *ndh* gene suite in photoautotrophs like the saguaro, and putatively other cacti.

The knowledge about plastome evolution across Opuntieae can help to elucidate the remarkable radiation of cactus diversity. *Opuntia* is the most species-rich lineage of the tribe, and likewise across all of Cactaceae (Korotkova *et al*., 2021). Despite this high diversity, which is considered to be quite young in origin (Arakaki *et al*. 2011), plastome sequences are shown to be extremely informative in recovering the phylogenetic relationships of its major clades. Additionally, the identification of different plastome features according to the presence or absence of the IR, content, and variation within the group promotes new insights into the diversification of the lineages and the putative drivers underlying such aspects, besides providing helpful signatures for barcoding and taxonomic assessment. For example, Chen *et al*. (2022) recently assembled a prickly pear plastome assigned to the *Opuntia sulphurea* taxon, which is a southern South American species. We analyzed their assembly in our dataset and detected it as a problematic identification based on plastome features. The plastome from Chen *et al*. (2022) is an IR-lacking sample (122 kb), much similar to the IR-lacking type 7 of our dataset (or the type 8, only present in *O. polyacantha*), typical of the North American *Opuntia* species that are not part of the Nopalea clade, while *Opuntia sulphurea* is part of the Elatae clade with which it shares their type 3 plastome features (Köhler *et al*., *unpublished data*). We confirm this by checking phylogenetic analyses of the GenBank accession MW927506 in our dataset using maximum-likelihood approaches with a reduced matrix utilizing the chloroplast markers listed by Köhler *et al*. (2020) as phylogenetic informative within Opuntioideae (Fig. S3), and suggest that the sample used in Chen *et al*. (2022) may represent *Opuntia polyacantha*, which slightly resembles *O. sulphurea* leading to erroneous identification.

Our phylogenetic analyses provided novel and robust relationships within Opuntieae lineages with a comprehensive sampling when compared to previous studies (Griffith & Porter, 2009; Majure *et al*., 2012; Majure & Puente, 2014; Köhler *et al*., 2020, 2021). However, in our study, we revealed that different datasets (whole plastome sequences vs. just genes) and alignment strategies (raw or trimmed with different allowances of gaps in Gblocks) have yielded distinct topologies involving mainly two nodes. Phylogenomic analyses are sensitive to systematic or alignment errors, and trimming biases (Zhong et al., 2011; Philippe et al., 2011; Philippe et al., 2017; Walker et al., 2019; Portik & Wiens, 2021), a pattern that was corroborated here. Furthermore, we demonstrated that few plastid markers can support a major supported topology (high bootstrap values), as well as underly contentious relationships. While some of these recalcitrant nodes can be the result of hard incongruences – with a set of different markers supporting alternative topologies – others can result from the lack of resolution across molecular sequences, usually leading to weak bootstrap support according to our analyses. Parins-Fukuchi *et al*. (2021) have demonstrated that phylogenomic conflicts coincide with rapid morphological innovations. The two contentious nodes recovered in our analyses may be one more representative of this phenomenon. The MSA clade represents a peculiar group within Opuntieae regarding morphology (with putative plesiomorphic characters, such as the terete stems in *Miqueliopuntia* and *Salmonopuntia*a) and geography (being endemic to southern South American arid regions of Atacama and inter-Andean valleys), which represents contrasting transitions to the other mostly ubiquitous characteristics of flattened stems and broader distribution of members of BT and *Opuntia* clades. Similarly, part of the Nopalea clade is characterized by members with unique floral features within Opuntieae showing hummingbird pollination syndrome, with short and erected tepals forming a tube with exserted stamens and styles. However, we are still lacking a nuclear phylogeny of the Opuntieae lineages to assess a more robust phylogenetic hypothesis, which will aid in understanding the evolutionary radiation of the group.

The issues involving topological incongruence among chloroplast markers have been recently explored (Gonçalves *et al*. 2019, 2020; Walker *et al*. 2019; Xiao *et al*., 2020; Zhang *et al*., 2020a,b), including within the cactus subfamily Opuntioideae (Köhler et a. 2020), and it has been suggested as a result of systematic errors that phylogenomic analyses are sensitive, or the product of biological events (e.g., heteroplasmy and horizontal gene transfer) which still need to be further investigated. In recent years, some studies have explored using the multispecies coalescent (MSC) approach for plastome markers stemming from the evidence of heteroplasmic recombination, however, most of the results tend to be confounding (Thode *et al*., 2021), while there are substantial arguments to continue treating plastomes as a single estimate of the underlying species phylogeny (Doyle, 2022). However, we submit that quantifying and filtering the phylogenetic signal in plastome data is an important step for evaluating topological concordance, especially when performing downstream comparative analyses based on this data.

## 5. CONCLUSIONS

Cacti are one of the most charismatic groups of plants, but with remarkable difficulties for taxonomic and systematic understanding due to several challenges such as hybridization, polyploidy, rapid diversification, the evolution of homoplasious characters, and the intimidating prickly morphology, which leads to a general lack of primary data, as few scientists feel comfortable putting their hands on them. In this study, we reinforce that more than a prickly morphology, cacti harbor intriguing molecular aspects related to their plastome variation putatively linked with their diversification history. Opuntieae, one impressive lineage of cacti, presents striking variation in plastome size related to contractions, expansions, and the loss of the inverted repeat region, as well as several cases of pseudogenization or gene loss. Despite some contentious signals across markers, analyses of incongruence can leverage plastome sequences to provide a robust framework to more deeply understand the evolutionary radiation of this group. Further studies putting our dataset into a broader scale across the cacti tree of life, or even exploring more closely related lineages, is still necessary. Besides, innovative analyses should be carried out to address how ecological drivers, physiological constraints, and morphological traits of cacti may be related to the variation in plastomes that have been reported across the family.

## ETHICS APPROVAL AND CONSENT TO PARTICIPATE

Not applicable.

## CONSENT FOR PUBLICATION

Not applicable.

## AVAILABILITY OF DATA AND MATERIALS

The dataset generated and analyzed in this study can be found in the GenBank submission GRP 8945163. Supplemental information cited is included in this published article.

## AUTHORS’ CONTRIBUTIONS

CRediT authorship statement. MK: Conceptualization, Methodology, Validation, Formal analysis, Investigation, Data Curation, Writing - original draft, Writing - review & editing, Visualization, Funding acquisition. MR: Methodology, Software, Validation, Formal analysis, Investigation, Writing - Review & Editing. JJ: Software, Methodology, Validation, Writing - review & editing. LCM: Conceptualization, Methodology, Validation, Investigation, Resources, Writing - review & editing, Funding acquisition. All authors read and approved the final manuscript.

## ACKNOWLEDGEMENTS

We thank the Desert Botanical Garden (DBG, Phoenix, AZ, USA) for some of the samples used in our analyses and for support for fieldwork in which some of the accessions used here were collected. MK is grateful to the American Society of Plant Taxonomists (ASPT), Cactus and Succulent Society of America (CSSA), International Association for Plant Taxonomy (IAPT), and IDEA WILD for financial support of part of the research here reported. MK also thanks the Brazilian National Council for Scientific and Technological Development (Conselho Nacional de Desenvolvimento Científico e Tecnológico – CNPq) for his PhD scholarship, the PDSE/CAPES (Process Number 88881.186882/2018-01) for supporting his period as Visiting Researcher at the Florida Museum of Natural History (FLMNH, UF, USA), and the PrInt/CAPES for granting his period as postdoctoral researcher (Process Number 88887.583192/2020-00). This study was also financed in part by the Coordenacão de Aperfeiçoamento de Pessoal de Nível Superior – Brasil (CAPES) – Finance Code 001, start-up funds to LCM from Florida Museum of Natural History and the University of Florida, and a National Science Foundation EAGER Grant (DEB #1735604) to Wojciechowski, Majure, Sanderson & Steele.

## FIGURE LEGENDS

**Figure S1.**
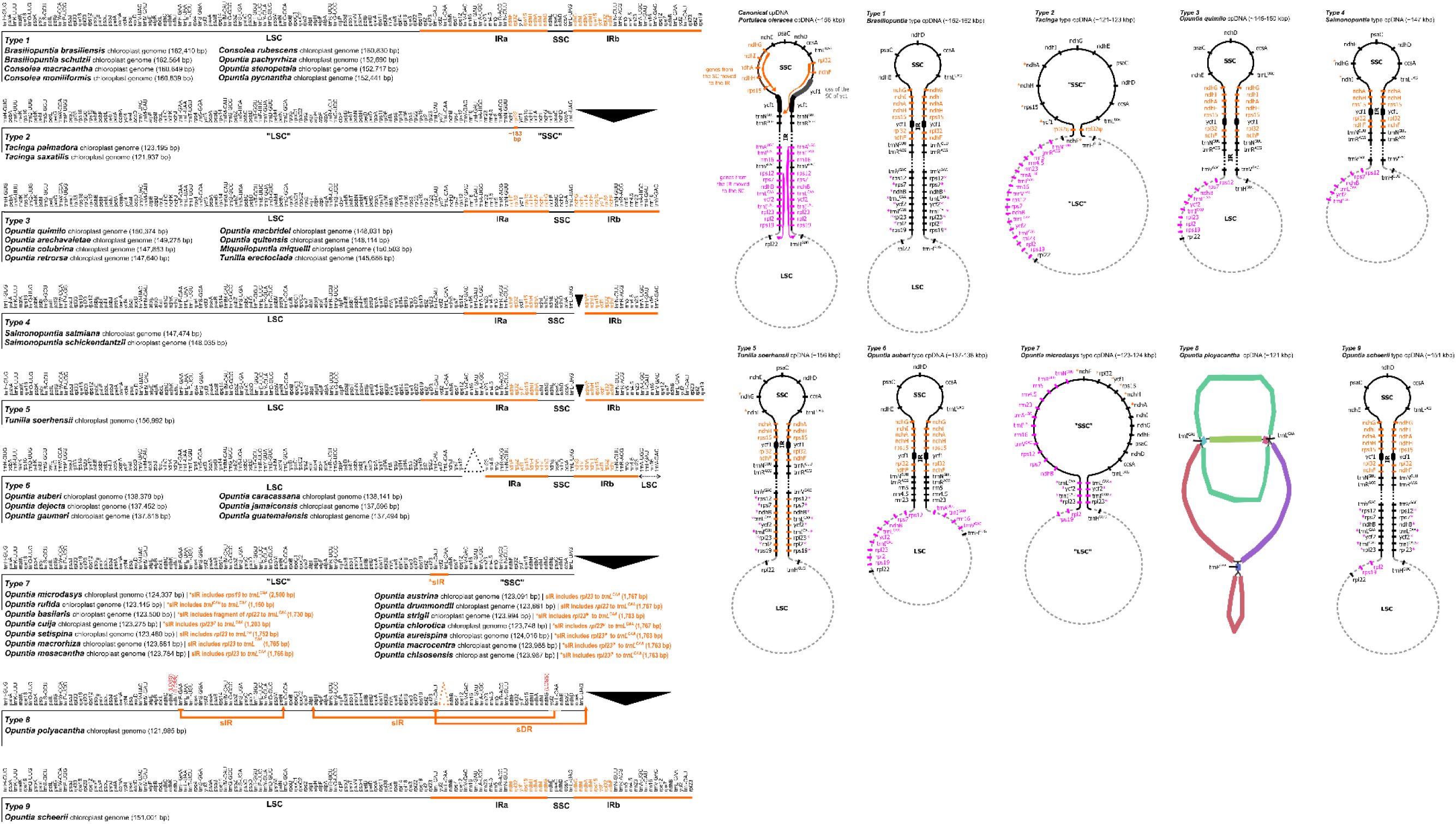
Type of plastomes assembled in Opuntieae samples. On the left, the circular genome map has been linearized, and the orange bars delimit the IR regions. Genes highlighted in orange are those incorporated into the IR, but typically found in the SSC of canonical plastomes. On the right, stereotypical plastome representation of circularized plastomes (except type 8). The arrows in orange highlight the movement of the genes typically found in the SSC to the IR regions (genes in orange), while the pink arrows highlight the movement of the genes typically found in the IR regions to the SC regions in Opuntieae plastomes.

**Figure S2.**
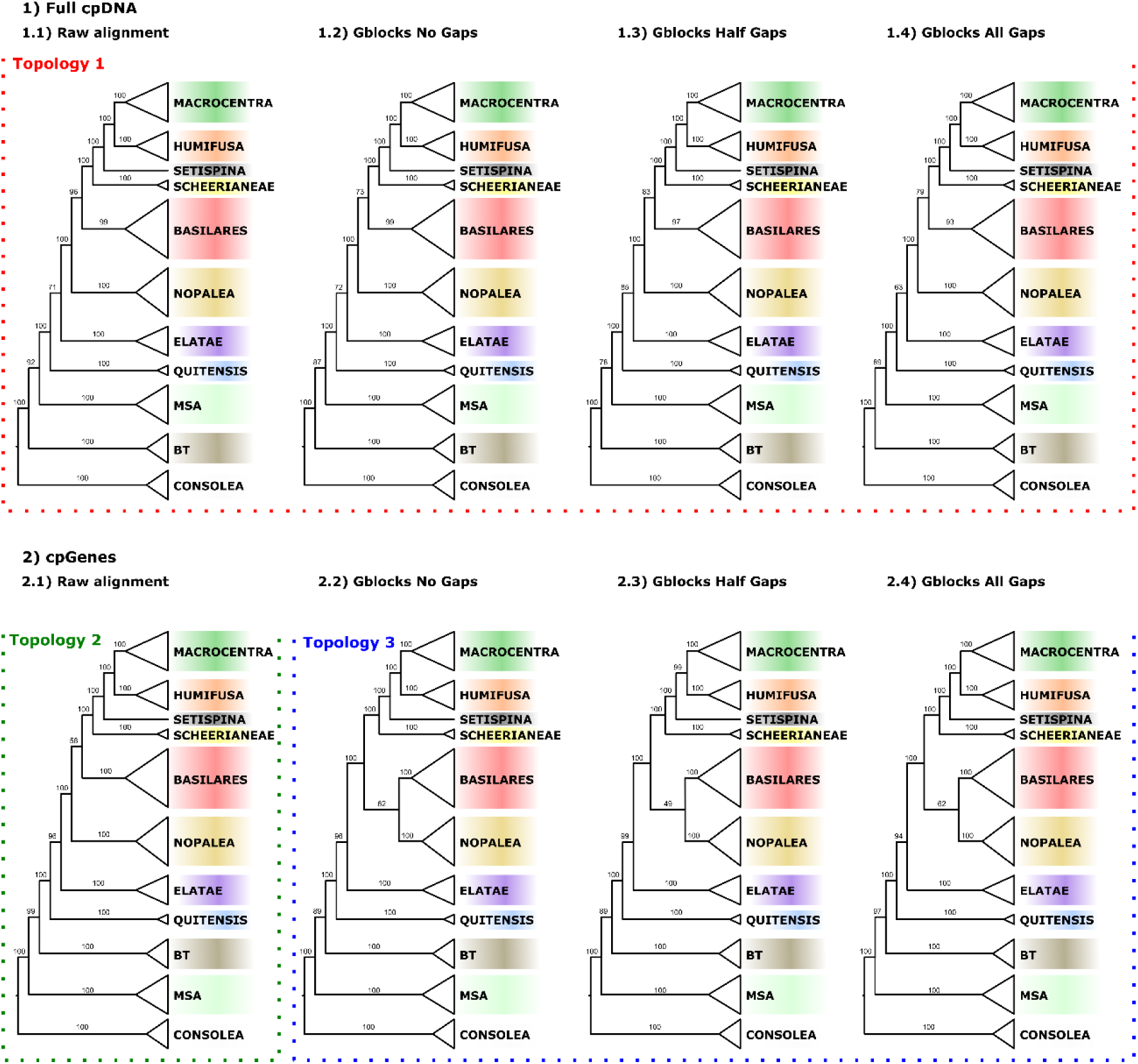
Maximum-likelihood inference of all data sets and alignment strategies analyzed. Full plastome (cpDNA) inferences recovered all the same major backcone topology. Plastidial genes (coding and non-coding regions) recovered different topologies from that of full plastome, and among alignment strategies (raw vs. trimmed with Gblocks).

**Figure S3.**
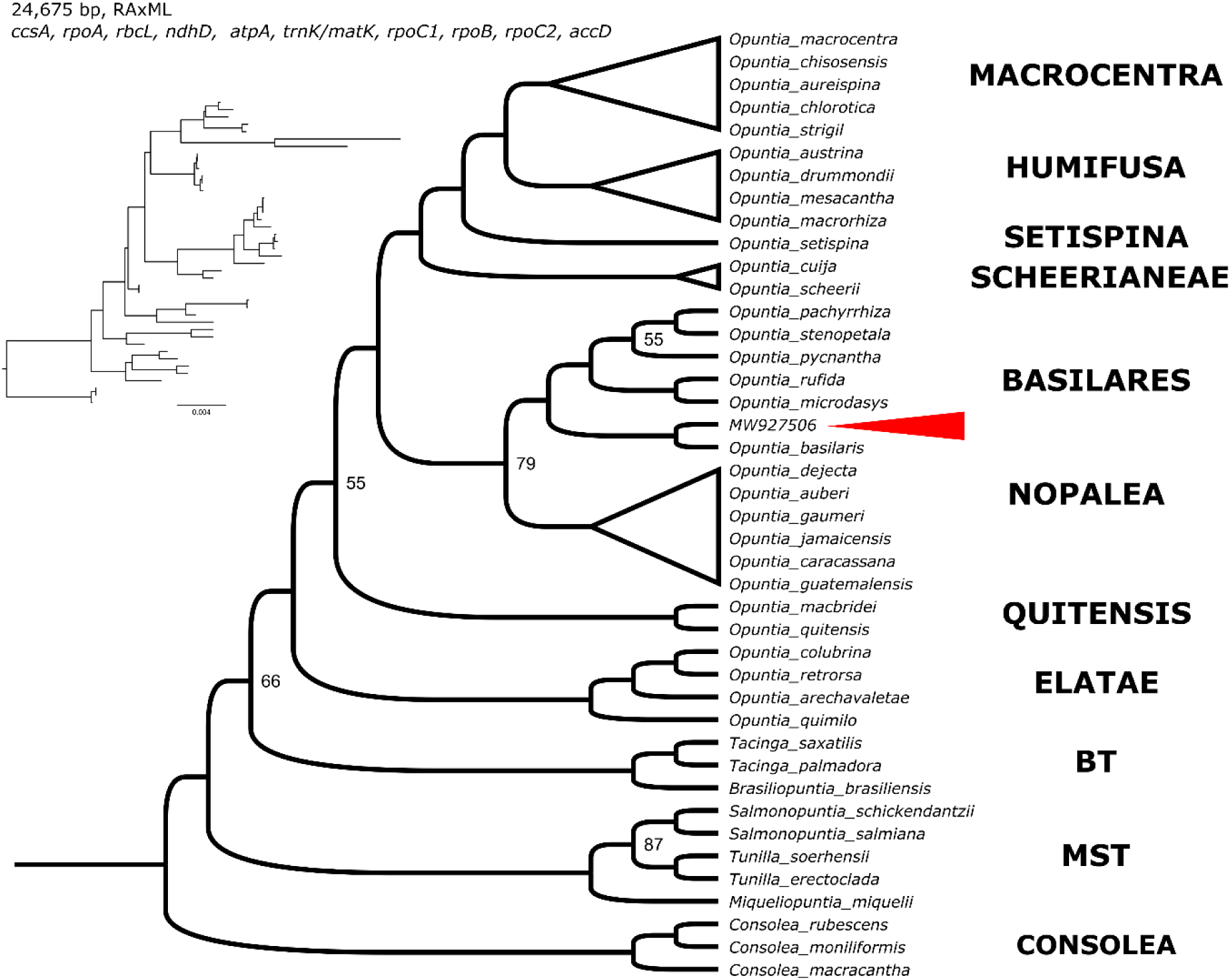
Phylogenetic placement of the GenBank accession MW927506 (from Chen *et al*., 2022) within the Basilares clade. The analysis is based on ten chloroplast markers listed by Köhler *et al*. (2020) as phylogenetic informative for Opuntioideae. Phylogenetic nodes have full bootstrap support (100), except when depicted. The accession MW927506, wrongly assigned as *O. sulphurea*, is highlighted with the red arrow, as part of the Basilares clade.

## TABLES

**Table S1.** Taxa sampled in this study, voucher information, raw read and post-quality control numbers, %GC content, and library content resulted from the chloroplast genome de novo assembly. Vouchers contain collector name, number and herbarium acronym in parentheses, or the accession number from the Desert Botanical Garden Living collection (https://dbg.org/research-conservation/living-collections/).

**Table S2.** Total number and variation (in length - bp) of tandem repeats identified in Opuntieae platomes.

**Table S3.** Statistics of alignments and maximum likelihood of each dataset.

**Table S4.** Complete gene-wise log-likelihood scores of topologies tests.

## REFERENCES

Abadi, S., Azouri, D., Pupko, T., Mayrose, I., 2019. Model selection may not be a mandatory step for phylogeny reconstruction. Nat Commun 10, 1–11. https://doi.org/10.1038/s41467-019-08822-w

Almeida, E.M., Sader, M.A., Rodriguez, P.E., Loeuille, B., Felix, L.P., Pedrosa-Harand, A., 2021. Assembling the puzzle: Complete chloroplast genome sequences of Discocactus bahiensis Britton & Rose and Melocactus ernestii Vaupel (Cactaceae) and their evolutionary significance. Braz. J. Bot 44, 877–888. https://doi.org/10.1007/s40415-021-00772-2

Amaral, D.T., Bombonato, J.R., da Silva Andrade, S.C., Moraes, E.M., Franco, F.F., 2021. The genome of a thorny species: comparative genomic analysis among South and North American Cactaceae. Planta 254, 44. https://doi.org/10.1007/s00425-021-03690-5

Anderson, E.F., 2001. The Cactus Family. Timber Press.

Arakaki, M., Christin, P.-A., Nyffeler, R., Lendel, A., Eggli, U., Ogburn, R.M., Spriggs, E., Moore, M.J., Edwards, E.J., 2011. Contemporaneous and recent radiations of the world’s major succulent plant lineages. PNAS 108, 8379–8384. https://doi.org/10.1073/pnas.1100628108

Bankevich, A., Nurk, S., Antipov, D., Gurevich, A.A., Dvorkin, M., Kulikov, A.S., Lesin, V.M., Nikolenko, S.I., Pham, S., Prjibelski, A.D., Pyshkin, A.V., Sirotkin, A.V., Vyahhi, N., Tesler, G., Alekseyev, M.A., Pevzner, P.A., 2012. SPAdes: A New Genome Assembly Algorithm and Its Applications to Single-Cell Sequencing. Journal of Computational Biology 19, 455–477. https://doi.org/10.1089/cmb.2012.0021

Becker, G., Lawrence, M., 2022. genbankr: Parsing GenBank files into semantically useful objects. R package version 1.27.0. [WWW Document]. URL https://bioconductor.org/packages/devel/bioc/html/genbankr.html

Bendich, A.J., 2004. Circular Chloroplast Chromosomes: The Grand Illusion. The Plant Cell 16, 1661–1666. https://doi.org/10.1105/tpc.160771

Bendich, A.J., Smith, S.B., 1990. Moving pictures and pulsed-field gel electrophoresis show linear DNA molecules from chloroplasts and mitochondria. Curr Genet 17, 421–425. https://doi.org/10.1007/BF00334522

Braukmann, T.W.A., Kuzmina, M., Stefanović, S., 2009. Loss of all plastid ndh genes in Gnetales and conifers: extent and evolutionary significance for the seed plant phylogeny. Curr Genet 55, 323–337. https://doi.org/10.1007/s00294-009-0249-7

Camacho, C., Coulouris, G., Avagyan, V., Ma, N., Papadopoulos, J., Bealer, K., Madden, T.L., 2009. BLAST+: architecture and applications. BMC Bioinformatics 10, 421. https://doi.org/10.1186/1471-2105-10-421

Castresana, J., 2000. Selection of Conserved Blocks from Multiple Alignments for Their Use in Phylogenetic Analysis. Molecular Biology and Evolution 17, 540–552. https://doi.org/10.1093/oxfordjournals.molbev.a026334

Cauz-Santos, L.A., da Costa, Z.P., Callot, C., Cauet, S., Zucchi, M.I., Bergès, H., van den Berg, C., Vieira, M.L.C., 2020. A Repertory of Rearrangements and the Loss of an Inverted Repeat Region in Passiflora Chloroplast Genomes. Genome Biology and Evolution 12, 1841–1857. https://doi.org/10.1093/gbe/evaa155

Chan, P.P., Lin, B.Y., Mak, A.J., Lowe, T.M., 2021. tRNAscan-SE 2.0: improved detection and functional classification of transfer RNA genes. Nucleic Acids Research 49, 9077–9096. https://doi.org/10.1093/nar/gkab688

Charboneau, J.L.M., Cronn, R.C., Liston, A., Wojciechowski, M.F., Sanderson, M.J., 2021. Plastome Structural Evolution and Homoplastic Inversions in Neo-Astragalus (Fabaceae). Genome Biology and Evolution 13, evab215. https://doi.org/10.1093/gbe/evab215

Chen, J., Zhang, S., Tang, W., Du, X., Yuan, Y., Wu, S., 2022. The complete chloroplast genome sequence of Opuntia sulphurea (Cactaceae). Mitochondrial DNA Part B 7, 361–362. https://doi.org/10.1080/23802359.2022.2035837

Choi, I.-S., Jansen, R., Ruhlman, T., 2020. Caught in the Act: Variation in plastid genome inverted repeat expansion within and between populations of Medicago minima. Ecology and Evolution 10, 12129–12137. https://doi.org/10.1002/ece3.6839

Cui, H., Ding, Z., Zhu, Q., Wu, Y., Qiu, B., Gao, P., 2021. Comparative analysis of nuclear, chloroplast, and mitochondrial genomes of watermelon and melon provides evidence of gene transfer. Sci Rep 11, 1595. https://doi.org/10.1038/s41598-020-80149-9

Dalla Costa, T.P., Silva, M.C., de Santana Lopes, A., Gomes Pacheco, T., de Oliveira, J.D., de Baura, V.A., Balsanelli, E., Maltempi de Souza, E., de Oliveira Pedrosa, F., Rogalski, M., 2022. The plastome of Melocactus glaucescens Buining & Brederoo reveals unique evolutionary features and loss of essential tRNA genes. Planta 255, 57. https://doi.org/10.1007/s00425-022-03841-2

Daniell, H., Lin, C.-S., Yu, M., Chang, W.-J., 2016. Chloroplast genomes: diversity, evolution, and applications in genetic engineering. Genome Biol 17, 134. https://doi.org/10.1186/s13059-016-1004-2

Darling, A.C.E., Mau, B., Blattner, F.R., Perna, N.T., 2004. Mauve: multiple alignment of conserved genomic sequence with rearrangements. Genome Res. 14, 1394–1403. https://doi.org/10.1101/gr.2289704

de Vries, J., Sousa, F.L., Bölter, B., Soll, J., Gould, S.B., 2015. YCF1: A Green TIC? Plant Cell 27, 1827–1833. https://doi.org/10.1105/tpc.114.135541

Downie, S.R., Palmer, J.D., 1994. A Chloroplast DNA Phylogeny of the Caryophyllales Based on Structural and Inverted Repeat Restriction Site Variation. Systematic Botany 19, 236–252. https://doi.org/10.2307/2419599

Doyle, J.J., 2022. Defining Coalescent Genes: Theory Meets Practice in Organelle Phylogenomics. Systematic Biology 71, 476–489. https://doi.org/10.1093/sysbio/syab053

Doyle, J.J., Doyle, J.L., 1987. A rapid DNA isolation procedure from small quantities of fresh leaf tissue. Phytochem. Bull. 19, 11–15.

Drescher, A., Ruf, S., Calsa, T., Carrer, H., Bock, R., 2000. The two largest chloroplast genome-encoded open reading frames of higher plants are essential genes. Plant J 22, 97–104. https://doi.org/10.1046/j.1365-313x.2000.00722.x

Dugas, D.V., Hernandez, D., Koenen, E.J.M., Schwarz, E., Straub, S., Hughes, C.E., Jansen, R.K., Nageswara-Rao, M., Staats, M., Trujillo, J.T., Hajrah, N.H., Alharbi, N.S., Al-Malki, A.L., Sabir, J.S.M., Bailey, C.D., 2015. Mimosoid legume plastome evolution: IR expansion, tandem repeat expansions and accelerated rate of evolution in clpP. Sci Rep 5, 16958. https://doi.org/10.1038/srep16958

Gitzendanner, M.A., Soltis, P.S., Wong, G.K.-S., Ruhfel, B.R., Soltis, D.E., 2018. Plastid phylogenomic analysis of green plants: A billion years of evolutionary history. American Journal of Botany 105, 291–301. https://doi.org/10.1002/ajb2.1048

Gonçalves, D.J.P., Jansen, R.K., Ruhlman, T.A., Mandel, J.R., 2020. Under the rug: Abandoning persistent misconceptions that obfuscate organelle evolution. Molecular Phylogenetics and Evolution 151, 106903. https://doi.org/10.1016/j.ympev.2020.106903

Gonçalves, D.J.P., Simpson, B.B., Ortiz, E.M., Shimizu, G.H., Jansen, R.K., 2019. Incongruence between gene trees and species trees and phylogenetic signal variation in plastid genes. Mol. Phylogenet. Evol. 138, 219–232. https://doi.org/10.1016/j.ympev.2019.05.022

Graham, S.W., Lam, V.K.Y., Merckx, V.S.F.T., 2017. Plastomes on the edge: the evolutionary breakdown of mycoheterotroph plastid genomes. New Phytologist 214, 48–55. https://doi.org/10.1111/nph.14398

Griffith, M.P., Porter, J.M., 2009. Phylogeny of Opuntioideae (Cactaceae). International Journal of Plant Sciences 170, 107–116. https://doi.org/10.1086/593048

Guisinger, M.M., Chumley, T.W., Kuehl, J.V., Boore, J.L., Jansen, R.K., 2010. Implications of the Plastid Genome Sequence of Typha (Typhaceae, Poales) for Understanding Genome Evolution in Poaceae. J Mol Evol 70, 149–166. https://doi.org/10.1007/s00239-009-9317-3

Harris, M.E., Meyer, G., Vandergon, T., Vandergon, V.O., 2013. Loss of the Acetyl-CoA Carboxylase (accD) Gene in Poales. Plant Mol Biol Rep 31, 21–31. https://doi.org/10.1007/s11105-012-0461-3

Jansen, R.K., Wojciechowski, M.F., Sanniyasi, E., Lee, S.-B., Daniell, H., 2008. Complete plastid genome sequence of the chickpea (Cicer arietinum) and the phylogenetic distribution of rps12 and clpP intron losses among legumes (Leguminosae). Molecular Phylogenetics and Evolution 48, 1204–1217. https://doi.org/10.1016/j.ympev.2008.06.013

Jin, D.-M., Wicke, S., Gan, L., Yang, J.-B., Jin, J.-J., Yi, T.-S., 2020a. The Loss of the Inverted Repeat in the Putranjivoid Clade of Malpighiales. Frontiers in Plant Science 11.

Jin, J.-J., Yu, W.-B., Yang, J.-B., Song, Y., de Pamphilis, C.W., Yi, T.-S., Li, D.-Z., 2020b. GetOrganelle: a fast and versatile toolkit for accurate de novo assembly of organelle genomes. Genome Biology 21, 241. https://doi.org/10.1186/s13059-020-02154-5

Jin, D.-M., Jin, J.-J., Yi, T.-S., 2020c. Plastome Structural Conservation and Evolution in the Clusioid Clade of Malpighiales. Sci Rep 10, 9091. https://doi.org/10.1038/s41598-020-66024-7

Jo, Y.D., Park, J., Kim, J., Song, W., Hur, C.-G., Lee, Y.-H., Kang, B.-C., 2011. Complete sequencing and comparative analyses of the pepper (Capsicum annuum L.) plastome revealed high frequency of tandem repeats and large insertion/deletions on pepper plastome. Plant Cell Rep 30, 217–229. https://doi.org/10.1007/s00299-010-0929-2

Johnson, M., 2017. PhypartsPieCharts [WWW Document]. URL https://github.com/mossmatters/phyloscripts/tree/master/phypartspiecharts (accessed 3.8.23).

Katoh, K., Standley, D.M., 2013. MAFFT multiple sequence alignment software version 7: improvements in performance and usability. Mol Biol Evol 30, 772–780. https://doi.org/10.1093/molbev/mst010

Kikuchi, S., Asakura, Y., Imai, M., Nakahira, Y., Kotani, Y., Hashiguchi, Y., Nakai, Y., Takafuji, K., Bédard, J., Hirabayashi-Ishioka, Y., Mori, H., Shiina, T., Nakai, M., 2018. A Ycf2-FtsHi Heteromeric AAA-ATPase Complex Is Required for Chloroplast Protein Import. The Plant Cell 30, 2677–2703. https://doi.org/10.1105/tpc.18.00357

Kim, H.T., Chase, M.W., 2017. Independent degradation in genes of the plastid ndh gene family in species of the orchid genus Cymbidium (Orchidaceae; Epidendroideae). PLOS ONE 12, e0187318. https://doi.org/10.1371/journal.pone.0187318

Köhler, M., Font, F., Puente-Martinez, R., Majure, L.C., 2021. “That’s Opuntia, that was!”, again: a new combination for an old and enigmatic Opuntia s.l. (Cactaceae). Phytotaxa 505, 262–274. https://doi.org/10.11646/phytotaxa.505.3.2

Köhler, M., Reginato, M., Souza-Chies, T.T., Majure, L.C., 2020. Insights Into Chloroplast Genome Evolution Across Opuntioideae (Cactaceae) Reveals Robust Yet Sometimes Conflicting Phylogenetic Topologies. Front. Plant Sci. 11. https://doi.org/10.3389/fpls.2020.00729

Korotkova, N., Aquino, D., Arias, S., Eggli, U., Franck, A., Gómez-Hinostrosa, C., Guerrero, P.C., Hernández, H.M., Kohlbecker, A., Köhler, M., Luther, K., Majure, L.C., Müller, A., Metzing, D., Nyffeler, R., Sánchez, D., Schlumpberger, B., Berendsohn, W.G., 2021. Cactaceae at Caryophyllales.org – a dynamic online species-level taxonomic backbone for the family. will 51, 251–270. https://doi.org/10.3372/wi.51.51208

Lee, C., Choi, I.-S., Cardoso, D., de Lima, H.C., de Queiroz, L.P., Wojciechowski, M.F., Jansen, R.K., Ruhlman, T.A., 2021. The chicken or the egg? Plastome evolution and an independent loss of the inverted repeat in papilionoid legumes. The Plant Journal 107, 861–875. https://doi.org/10.1111/tpj.15351

Li, F.-W., Kuo, L.-Y., Pryer, K.M., Rothfels, C.J., 2016. Genes Translocated into the Plastid Inverted Repeat Show Decelerated Substitution Rates and Elevated GC Content. Genome Biology and Evolution 8, 2452–2458. https://doi.org/10.1093/gbe/evw167

Lilly, J.W., Havey, M.J., Jackson, S.A., Jiang, J., 2001. Cytogenomic Analyses Reveal the Structural Plasticity of the Chloroplast Genome in Higher Plants. The Plant Cell 13, 245–254. https://doi.org/10.1105/tpc.13.2.245

Lin, C.-S., Chen, J.J.W., Chiu, C.-C., Hsiao, H.C.W., Yang, C.-J., Jin, X.-H., Leebens-Mack, J., de Pamphilis, C.W., Huang, Y.-T., Yang, L.-H., Chang, W.-J., Kui, L., Wong, G.K.-S., Hu, J.-M., Wang, W., Shih, M.-C., 2017. Concomitant loss of NDH complex-related genes within chloroplast and nuclear genomes in some orchids. The Plant Journal 90, 994–1006. https://doi.org/10.1111/tpj.13525

Lin, C.-S., Chen, J.J.W., Huang, Y.-T., Chan, M.-T., Daniell, H., Chang, W.-J., Hsu, C.-T., Liao, D.-C., Wu, F.-H., Lin, S.-Y., Liao, C.-F., Deyholos, M.K., Wong, G.K.-S., Albert, V.A., Chou, M.-L., Chen, C.-Y., Shih, M.-C., 2015. The location and translocation of ndh genes of chloroplast origin in the Orchidaceae family. Sci Rep 5, 9040. https://doi.org/10.1038/srep09040

Liu, X., Yang, H., Zhao, J., Zhou, B., Li, T., Xiang, B., 2018. The complete chloroplast genome sequence of the folk medicinal and vegetable plant purslane (Portulaca oleracea L.). The Journal of Horticultural Science and Biotechnology 93, 356–365. https://doi.org/10.1080/14620316.2017.1389308

Majure, L.C., Baker, M.A., Cloud-Hughes, M., Salywon, A., Neubig, K.M., 2019. Phylogenomics in Cactaceae: A case study using the chollas sensu lato (Cylindropuntieae, Opuntioideae) reveals a common pattern out of the Chihuahuan and Sonoran deserts. Am. J. Bot. 106, 1327–1345. https://doi.org/10.1002/ajb2.1364

Majure, L.C., Puente, R., 2014. Phylogenetic relationships and morphological evolution in Opuntia s. str. and closely related members of tribe Opuntieae. Succulent Plant Research 8, 9–30.

Majure, L.C., Puente, R., Griffith, M.P., Judd, W.S., Soltis, P.S., Soltis, D.E., 2012. Phylogeny of Opuntia s.s. (Cactaceae): clade delineation, geographic origins, and reticulate evolution. Am. J. Bot. 99, 847–864. https://doi.org/10.3732/ajb.1100375

Mayer, C., 2006. Phobos 3.3. 11 (http://www.rub.de/spezzoo/cm/cm_phobos.htm).

Miller, M.A., Pfeiffer, W., Schwartz, T., 2010. Creating the CIPRES Science Gateway for inference of large phylogenetic trees, in: 2010 Gateway Computing Environments Workshop (GCE). Presented at the 2010 Gateway Computing Environments Workshop (GCE), pp. 1–8. https://doi.org/10.1109/GCE.2010.5676129

Mower, J.P., Vickrey, T.L., 2018. Structural Diversity Among Plastid Genomes of Land Plants, in: Chaw, S.-M., Jansen, R.K. (Eds.), Advances in Botanical Research, Plastid Genome Evolution. Academic Press, pp. 263–292. https://doi.org/10.1016/bs.abr.2017.11.013

Oldenburg, D.J., Bendich, A.J., 2016. The linear plastid chromosomes of maize: terminal sequences, structures, and implications for DNA replication. Curr Genet 62, 431–442. https://doi.org/10.1007/s00294-015-0548-0

Oldenburg, D.J., Bendich, A.J., 2015. DNA maintenance in plastids and mitochondria of plants. Frontiers in Plant Science 6.

Oldenburg, D.J., Bendich, A.J., 2004. Most Chloroplast DNA of Maize Seedlings in Linear Molecules with Defined Ends and Branched Forms. Journal of Molecular Biology 335, 953– 970. https://doi.org/10.1016/j.jmb.2003.11.020

Palmer, J.D., Jansen, R.K., Michaels, H.J., Chase, M.W., Manhart, J.R., 1988. Chloroplast DNA Variation and Plant Phylogeny. Annals of the Missouri Botanical Garden 75, 1180–1206. https://doi.org/10.2307/2399279

Palmer, J.D., Thompson, W.F., 1982. Chloroplast DNA rearrangements are more frequent when a large inverted repeat sequence is lost. Cell 29, 537–550. https://doi.org/10.1016/0092-8674(82)90170-2

Paradis, E., Schliep, K., 2019. ape 5.0: an environment for modern phylogenetics and evolutionary analyses in R. Bioinformatics 35, 526–528. https://doi.org/10.1093/bioinformatics/bty633

Parins-Fukuchi, C., Stull, G.W., Smith, S.A., 2021. Phylogenomic conflict coincides with rapid morphological innovation. Proceedings of the National Academy of Sciences 118, e2023058118. https://doi.org/10.1073/pnas.2023058118

Peredo, E.L., King, U.M., Les, D.H., 2013. The Plastid Genome of Najas flexilis: Adaptation to Submersed Environments Is Accompanied by the Complete Loss of the NDH Complex in an Aquatic Angiosperm. PLOS ONE 8, e68591. https://doi.org/10.1371/journal.pone.0068591

Philippe, H., Brinkmann, H., Lavrov, D.V., Littlewood, D.T.J., Manuel, M., Wörheide, G., Baurain, D., 2011. Resolving Difficult Phylogenetic Questions: Why More Sequences Are Not Enough. PLOS Biology 9, e1000602. https://doi.org/10.1371/journal.pbio.1000602

Philippe, H., Vienne, D.M. de Ranwez, V., Roure, B., Baurain, D., Delsuc, F., 2017. Pitfalls in supermatrix phylogenomics. European Journal of Taxonomy. https://doi.org/10.5852/ejt.2017.283

Portik, D.M., Wiens, J.J., 2021. Do Alignment and Trimming Methods Matter for Phylogenomic (UCE) Analyses? Systematic Biology 70, 440–462. https://doi.org/10.1093/sysbio/syaa064

Qin, Q., Li, J., Zeng, S., Xu, Y., Han, F., Yu, J., 2022. The complete plastomes of red fleshed pitaya (Selenicereus monacanthus) and three related Selenicereus species: insights into gene losses, inverted repeat expansions and phylogenomic implications. Physiol Mol Biol Plants 28, 123–137. https://doi.org/10.1007/s12298-021-01121-z

Rabah, S.O., Shrestha, B., Hajrah, N.H., Sabir, Mumdooh J., Alharby, H.F., Sabir, Mernan J., Alhebshi, A.M., Sabir, J.S.M., Gilbert, L.E., Ruhlman, T.A., Jansen, R.K., 2019. Passiflora plastome sequencing reveals widespread genomic rearrangements. Journal of Systematics and Evolution 57, 1–14. https://doi.org/10.1111/jse.12425

Raubeson, L.A., Jansen, R.K., 2005. Chloroplast genomes of plants., in: Henry, R.J. (Ed.), Plant Diversity and Evolution: Genotypic and Phenotypic Variation in Higher Plants. CABI, Wallingford, pp. 45–68. https://doi.org/10.1079/9780851999043.0045

Ruhlman, T.A., Chang, W.-J., Chen, J.J., Huang, Y.-T., Chan, M.-T., Zhang, J., Liao, D.-C., Blazier, J.C., Jin, X., Shih, M.-C., Jansen, R.K., Lin, C.-S., 2015. NDH expression marks major transitions in plant evolution and reveals coordinate intracellular gene loss. BMC Plant Biology 15, 100. https://doi.org/10.1186/s12870-015-0484-7

Ruhlman, T.A., Jansen, R.K., 2021. Plastid Genomes of Flowering Plants: Essential Principles, in: Maliga, P. (Ed.), Chloroplast Biotechnology: Methods and Protocols, Methods in Molecular Biology. Springer US, New York, NY, pp. 3–47. https://doi.org/10.1007/978-1-0716-1472-3_1

Ruhlman, T.A., Jansen, R.K., 2018. Aberration or Analogy? The Atypical Plastomes of Geraniaceae, in: Chaw, S.-M., Jansen, R.K. (Eds.), Advances in Botanical Research, Plastid Genome Evolution. Academic Press, pp. 223–262. https://doi.org/10.1016/bs.abr.2017.11.017

Sadali, N.M., Sowden, R.G., Ling, Q., Jarvis, R.P., 2019. Differentiation of chromoplasts and other plastids in plants. Plant Cell Rep 38, 803–818. https://doi.org/10.1007/s00299-019-02420-2

Sanderson, M.J., Copetti, D., Búrquez, A., Bustamante, E., Charboneau, J.L.M., Eguiarte, L.E., Kumar, S., Lee, H.O., Lee, J., McMahon, M., Steele, K., Wing, R., Yang, T.-J., Zwickl, D., Wojciechowski, M.F., 2015. Exceptional reduction of the plastid genome of saguaro cactus (Carnegiea gigantea): Loss of the ndh gene suite and inverted repeat. Am J Bot 102, 1115–1127. https://doi.org/10.3732/ajb.1500184

Schliep, K.P., 2011. phangorn: phylogenetic analysis in R. Bioinformatics 27, 592–593. https://doi.org/10.1093/bioinformatics/btq706

Shen, X.-X., Hittinger, C.T., Rokas, A., 2017. Contentious relationships in phylogenomic studies can be driven by a handful of genes. Nat Ecol Evol 1, 1–10. https://doi.org/10.1038/s41559-017-0126

Shimodaira, H., Hasegawa, M., 1999. Multiple Comparisons of Log-Likelihoods with Applications to Phylogenetic Inference. Molecular Biology and Evolution 16, 1114. https://doi.org/10.1093/oxfordjournals.molbev.a026201

Silva, G.M. da de Santana Lopes, A., Gomes Pacheco, T., Lima de Godoy Machado, K., Silva, M.C., de Oliveira, J.D., de Baura, V.A., Balsanelli, E., Maltempi de Souza, E., de Oliveira Pedrosa, F., Rogalski, M., 2021. Genetic and evolutionary analyses of plastomes of the subfamily Cactoideae (Cactaceae) indicate relaxed protein biosynthesis and tRNA import from cytosol. Braz. J. Bot 44, 97–116. https://doi.org/10.1007/s40415-020-00689-2

Silva, S.R., Diaz, Y.C.A., Penha, H.A., Pinheiro, D.G., Fernandes, C.C., Miranda, V.F.O., Michael, T.P., Varani, A.M., 2016. The Chloroplast Genome of Utricularia reniformis Sheds Light on the Evolution of the ndh Gene Complex of Terrestrial Carnivorous Plants from the Lentibulariaceae Family. PLOS ONE 11, e0165176. https://doi.org/10.1371/journal.pone.0165176

Sinn, B.T., Sedmak, D.D., Kelly, L.M., Freudenstein, J.V., 2018. Total duplication of the small single copy region in the angiosperm plastome: Rearrangement and inverted repeat instability in Asarum. American Journal of Botany 105, 71–84. https://doi.org/10.1002/ajb2.1001

Smith, S.A., Moore, M.J., Brown, J.W., Yang, Y., 2015. Analysis of phylogenomic datasets reveals conflict, concordance, and gene duplications with examples from animals and plants. BMC Evolutionary Biology 15, 150. https://doi.org/10.1186/s12862-015-0423-0

Solórzano, S., Chincoya, D.A., Sanchez-Flores, A., Estrada, K., Díaz-Velásquez, C.E., González-Rodríguez, A., Vaca-Paniagua, F., Dávila, P., Arias, S., 2019. De Novo Assembly Discovered Novel Structures in Genome of Plastids and Revealed Divergent Inverted Repeats in Mammillaria (Cactaceae, Caryophyllales). Plants (Basel) 8. https://doi.org/10.3390/plants8100392

Stamatakis, A., 2014. RAxML version 8: a tool for phylogenetic analysis and post-analysis of large phylogenies. Bioinformatics 30, 1312–1313. https://doi.org/10.1093/bioinformatics/btu033

Stegemann, S., Hartmann, S., Ruf, S., Bock, R., 2003. High-frequency gene transfer from the chloroplast genome to the nucleus. Proceedings of the National Academy of Sciences 100, 8828–8833. https://doi.org/10.1073/pnas.1430924100

Strand, D.D., D’Andrea, L., Bock, R., 2019. The plastid NAD(P)H dehydrogenase-like complex: structure, function and evolutionary dynamics. Biochemical Journal 476, 2743–2756. https://doi.org/10.1042/BCJ20190365

Straub, S.C.K., Parks, M., Weitemier, K., Fishbein, M., Cronn, R.C., Liston, A., 2012. Navigating the tip of the genomic iceberg: Next-generation sequencing for plant systematics. American Journal of Botany 99, 349–364. https://doi.org/10.3732/ajb.1100335

Talavera, G., Castresana, J., 2007. Improvement of Phylogenies after Removing Divergent and Ambiguously Aligned Blocks from Protein Sequence Alignments. Systematic Biology 56, 564–577. https://doi.org/10.1080/10635150701472164

Thode, V.A., Oliveira, C.T., Loeuille, B., Siniscalchi, C.M., Pirani, J.R., 2021. Comparative analyses of Mikania (Asteraceae: Eupatorieae) plastomes and impact of data partitioning and inference methods on phylogenetic relationships. Sci Rep 11, 13267. https://doi.org/10.1038/s41598-021-92727-6

Tillich, M., Lehwark, P., Pellizzer, T., Ulbricht-Jones, E.S., Fischer, A., Bock, R., Greiner, S., 2017. GeSeq - versatile and accurate annotation of organelle genomes. Nucleic Acids Res. 45, W6– W11. https://doi.org/10.1093/nar/gkx391

Timmis, J.N., Ayliffe, M.A., Huang, C.Y., Martin, W., 2004. Endosymbiotic gene transfer: organelle genomes forge eukaryotic chromosomes. Nat Rev Genet 5, 123–135. https://doi.org/10.1038/nrg1271

Walker, J.F., Jansen, R.K., Zanis, M.J., Emery, N.C., 2015. Sources of inversion variation in the small single copy (SSC) region of chloroplast genomes. American Journal of Botany 102, 1751–1752. https://doi.org/10.3732/ajb.1500299

Walker, J.F., Walker-Hale, N., Vargas, O.M., Larson, D.A., Stull, G.W., 2019. Characterizing gene tree conflict in plastome-inferred phylogenies. PeerJ 7, e7747. https://doi.org/10.7717/peerj.7747

Walker, J.F., Yang, Y., Feng, T., Timoneda, A., Mikenas, J., Hutchison, V., Edwards, C., Wang, N., Ahluwalia, S., Olivieri, J., Walker-Hale, N., Majure, L.C., Puente, R., Kadereit, G., Lauterbach, M., Eggli, U., Flores-Olvera, H., Ochoterena, H., Brockington, S.F., Moore, M.J., Smith, S.A., 2018. From cacti to carnivores: Improved phylotranscriptomic sampling and hierarchical homology inference provide further insight into the evolution of Caryophyllales. American Journal of Botany 105, 446–462. https://doi.org/10.1002/ajb2.1069

Wallace, R.S., 1995. Molecular systematic study of the Cactaceae: Using chloroplast DNA variation to elucidate Cactus phylogeny. brad 1995, 1–12. https://doi.org/10.25223/brad.n13.1995.a1

Wang, W.-C., Chen, S.-Y., Zhang, X.-Z., 2016. Chloroplast Genome Evolution in Actinidiaceae: clpP Loss, Heterogenous Divergence and Phylogenomic Practice. PLOS ONE 11, e0162324. https://doi.org/10.1371/journal.pone.0162324

Wick, R.R., Schultz, M.B., Zobel, J., Holt, K.E., 2015. Bandage: interactive visualization of de novo genome assemblies. Bioinformatics 31, 3350–3352. https://doi.org/10.1093/bioinformatics/btv383

Wicke, S., Schneeweiss, G.M., de Pamphilis, C.W., Müller, K.F., Quandt, D., 2011. The evolution of the plastid chromosome in land plants: gene content, gene order, gene function. Plant Mol Biol 76, 273–297. https://doi.org/10.1007/s11103-011-9762-4

Wu, S., Chen, J., Li, Y., Liu, A., Li, A., Yin, M., Shrestha, N., Liu, J., Ren, G., 2021. Extensive genomic rearrangements mediated by repetitive sequences in plastomes of Medicago and its relatives. BMC Plant Biol 21, 421. https://doi.org/10.1186/s12870-021-03202-3

Xiao, T.-W., Xu, Y., Jin, L., Liu, T.-J., Yan, H.-F., Ge, X.-J., 2020. Conflicting phylogenetic signals in plastomes of the tribe Laureae (Lauraceae). PeerJ 8. https://doi.org/10.7717/peerj.10155

Xu, X., Wang, D., 2021. Comparative Chloroplast Genomics of Corydalis Species (Papaveraceae): Evolutionary Perspectives on Their Unusual Large Scale Rearrangements. Frontiers in Plant Science 11.

Yang, X.-F., Wang, Y.-T., Chen, S.-T., Li, J.-K., Shen, H.-T., Guo, F.-Q., 2016. PBR1 selectively controls biogenesis of photosynthetic complexes by modulating translation of the large chloroplast gene Ycf1 in Arabidopsis. Cell Discov 2, 16003. https://doi.org/10.1038/celldisc.2016.3

Yao, G., Jin, J.-J., Li, H.-T., Yang, J.-B., Mandala, V.S., Croley, M., Mostow, R., Douglas, N.A., Chase, M.W., Christenhusz, M.J.M., Soltis, D.E., Soltis, P.S., Smith, S.A., Brockington, S.F., Moore, M.J., Yi, T.-S., Li, D.-Z., 2019. Plastid phylogenomic insights into the evolution of Caryophyllales. Molecular Phylogenetics and Evolution 134, 74–86. https://doi.org/10.1016/j.ympev.2018.12.023

Yu, J., Li, J., Zuo, Y., Qin, Q., Zeng, S., Rennenberg, H., Deng, H., 2023. Plastome variations reveal the distinct evolutionary scenarios of plastomes in the subfamily Cereoideae (Cactaceae). BMC Plant Biology 23, 132. https://doi.org/10.1186/s12870-023-04148-4

Zhang, R., Wang, Y.-H., Jin, J.-J., Stull, G.W., Bruneau, A., Cardoso, D., De Queiroz, L.P., Moore, M.J., Zhang, S.-D., Chen, S.-Y., Wang, J., Li, D.-Z., Yi, T.-S., 2020a. Exploration of Plastid Phylogenomic Conflict Yields New Insights into the Deep Relationships of Leguminosae. Systematic Biology 69, 613–622. https://doi.org/10.1093/sysbio/syaa013

Zhang, X., Sun, Y., Landis, J.B., Lv, Z., Shen, J., Zhang, H., Lin, N., Li, L., Sun, J., Deng, T., Sun, H., Wang, H., 2020b. Plastome phylogenomic study of Gentianeae (Gentianaceae): widespread gene tree discordance and its association with evolutionary rate heterogeneity of plastid genes. BMC Plant Biology 20, 340. https://doi.org/10.1186/s12870-020-02518-w

Zhong, B., Deusch, O., Goremykin, V.V., Penny, D., Biggs, P.J., Atherton, R.A., Nikiforova, S.V., Lockhart, P.J., 2011. Systematic Error in Seed Plant Phylogenomics. Genome Biology and Evolution 3, 1340–1348. https://doi.org/10.1093/gbe/evr105

Zhu, A., Guo, W., Gupta, S., Fan, W., Mower, J.P., 2016. Evolutionary dynamics of the plastid inverted repeat: the effects of expansion, contraction, and loss on substitution rates. New Phytologist 209, 1747–1756. https://doi.org/10.1111/nph.13743

